# Resolving the Activation Mechanism of the Human 20S Proteasome

**DOI:** 10.64898/2026.04.13.718244

**Authors:** Bryan D. Ryder, Nicholas L. Yan, Danna Trejos-Vidal, Patricia Martínez-Botía, Julian R. Braxton, Andrew Lim, Hannah Felstead, Steve Andrews, Eric Tse, Arthur A. De Melo, John Skidmore, Miguel A. Prado, Daniel R. Southworth, Jason E. Gestwicki

## Abstract

Proteasome activators (PAs) bind the α-subunits of the human 20S proteasome (h20S), opening the “gates” and allowing entry of substrate proteins. Aging is associated with diminished proteasome activity, leading to interest in understanding this activation mechanism. Evolving models have been proposed regarding PAs C-terminal tails, yet the critical molecular contacts for gate-opening are unclear. Here, we show a conserved leucine in the 5th position (P5) of the C-terminus is essential for h20S gate opening. By engineering C-termini in a model activator, PA26_E102A_, we show mutations to P5 systematically modulate proteasome activity *in vitro* and in cells. Structures of PA26_E102A_:h20S complexes at 2.7-3.2 Å resolution identify interactions between P5 and a conserved arginine in the h20S, leading to partial or full gate opening. These results clarify the essential contacts required for h20S gate opening, potentially enabling design of proteasome activation therapies.

## INTRODUCTION

The ubiquitin proteasome system (UPS) and the ubiquitin-independent proteasome system (UIPS) are major pathways for protein turnover^1–3^. The key complex in both systems is the 20S proteasome (20S); a homodimer of two hetero-heptameric rings that encircle a central channel. Substrate entry into the channel is regulated by the outer α-subunit rings, while catalytic protease activity is governed by the inner β-subunit rings^4,5^ (**Fig. 1a**). In the closed state, the N-terminal “gates” of the α-subunits obstruct the axial center channel and prevent most substrates from entering. Upon binding to a proteasome activator (PA), the N-termini of the 20S α-subunits retract, allowing protein substrates to enter the catalytic chamber^6–9^. This critical activation process is termed “gate opening”, which is required for substrate degradation.

**Figure 1:**
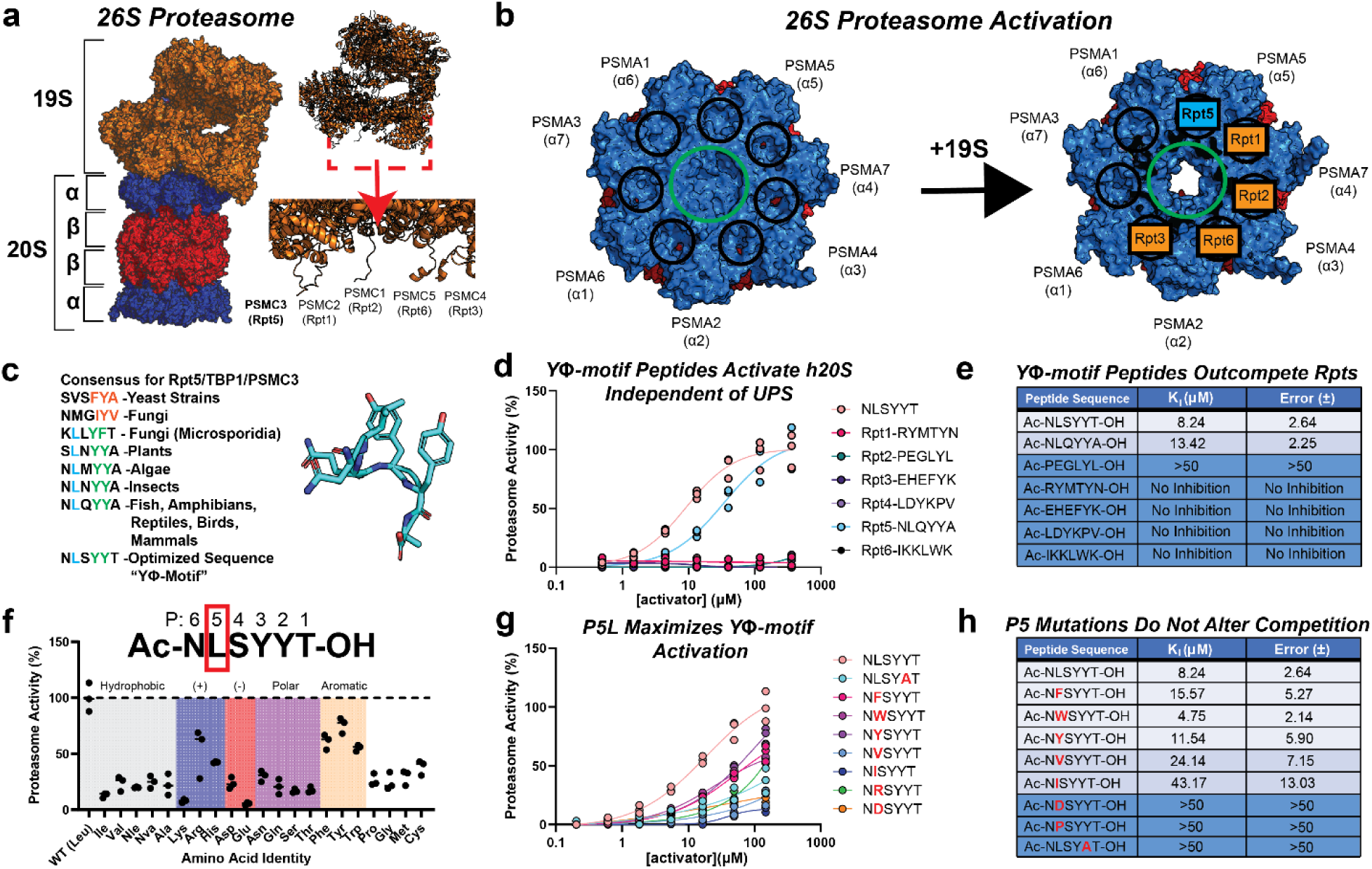
Proteasome Activation Relies on Sequence Conserved Leucine. **a)** Structural overview of the human 26S Proteasome (PDBID: 6MSK)^23^. The 26S complex contains the 19S proteasome activator (orange) and the 20S proteasome. The 20S contains regulatory α-subunits (blue) and catalytic β-subunits (red). Subunits are labeled with their human gene (PSMC*) and yeast homolog (Rpt*) names. **b)** Top view of the 20S with activation pockets (black circles) and the central gate channel (green circle) highlighted in the closed (left) and open (right) states. All α-subunits are labeled with human gene (PSMA*) and yeast homolog (α*) names. 19S subunits are shown over their corresponding binding pockets (orange boxes) with the Rpt5 subunit highlighted (cyan box). **c)** Rpt5 C-terminal sequences across different eukaryotic organisms. Tails containing HbYX motifs are marked in red, while tails containing YФ are marked in green and co-evolving P5L in blue. The structure of PA optimized peptide NLSYYT is shown to the right (PDBID: 6XMJ)^25^. **d)** Proteasome chymotrypsin-like activity assay. Maximum activity was normalized to NLSYYT. NLSYYT peptide (salmon, EC_50_ = 9.18 ± 2.25 μM) and Rpt5 (pink, EC_50_ = 34.02 ± 15.51 μM) were the only peptides to stimulate proteasome activity independently. Data are plotted individually (n=3). **e)** Table of estimated K_I_ values based on a fluorescence polarization competition assay. Reported values are averaged (n=3). **f)** Chymotrypsin-like activity assay where the 5^th^ C-terminal amino acid in each peptide (P5, red box) was replaced by all listed amino acids that are small hydrophobic (grey), basic (blue), acidic (red), polar (purple), aromatic (beige), or other (white). All peptides were normalized to NLSYYT (100 ± 12.77%) with Phe (61.01 ± 6.75%), Trp (55.32 ± 2.86%), and Tyr (75.78 ± 6.97%) showing reduced activity and all others showing significant loss of activity (<50%). Data are plotted individually (n=3). **g)** Proteasome chymotrypsin-like activity assay with peptides derived from NLSYYT with P5 substitutions (red letters). Maximum activity was normalized to NLSYYT. Average curve fits are shown with data plotted individually (n=3). NLSYYT (salmon, EC_50_ = 18.36 ± 8.78 μM), NFSYYT (pink, EC_50_ = 52.54 ± 32.09 μM), NWSYYT (lavender, EC_50_ = 24.91 ± 3.44 μM), and NYSYYT (purple, EC_50_ = 103.8 ± 12.87 μM) are the only four peptides to show significant activity. **h)** Table of estimated K_I_ values from a fluorescence polarization competition assay with peptides derived from NLSYYT with P5 substitutions (red letters). Reported values are averaged (n=3).

Proteasome activity is known to diminish during aging^2,3^. In some cell types, this decreased activity is a product of the relative accumulation of “closed” h20S particles and a decreased abundance of active, “open” UPS particles^2,8^. Also, the substrates of the UIPS, such as intrinsically disordered proteins^10,11^, accumulate with aging, suggesting that this system becomes less active. These observations have created interest in the development of anti-aging approaches that promote proteasome gate opening and activity^3,11^. For example, it has been shown that treatment with PA-derived peptides facilitates degradation of disease-associated, intrinsically disordered proteins (IDPs)^9,10^. Further, genetically deleting the 20S proteasome gates has been shown to accelerate turnover of some IDPs^12,13^ and promote longevity in worms^14^. We propose that these translational efforts would benefit from a deeper molecular understanding of how the natural PAs work.

Pioneering studies in model organisms have explored the mechanisms of gate opening. One prominent mechanism is mediated by the C-terminal tails of PAs, which insert into pockets at the protomer interface between α-subunits^5,6,15^. Studies on archaeal and yeast PAs showed that most C-terminal sequences end in a HbYX motif consisting of a hydrophobic (Hb) amino acid, tyrosine (Y), and any amino acid (X) as the three most C-terminal residues^1,6^. Part of this mechanism appears to be evolutionarily conserved because the HbYX motif is found in other eukaryotic PAs, including human PA200 and 3 of the 6 subunits of the human 19S complex: PSMC1, PSMC3, and PSMC4^1,16–20^ (**Fig. 1a**). Structural studies in the archaeal system^18,21^ advanced this knowledge by revealing how hydrogen bonding (H-bond) and hydrophobic contacts dock the HbYX motif into the α-subunit pockets. Because of this strong history, naming conventions sometimes refer to the human proteasome subunits by their yeast homolog, with human 19S “PSMC” subunits equivalent to the *S. cerevisae* “Rpt” subunits^9^ and h20S core particle subunits named by the broader yeast terms: α and β. Throughout this manuscript, we primarily adopt this strategy, using the yeast homolog nomenclature to describe the subunits of the 20S proteasome and 19S activator (**Fig. 1a,b**).

While extensive studies have defined PA engagement in model organisms, the activation mechanism of the human 20S (h20S) proteasome appears to be more complex. For example, structural studies^15,17,18,20^ showed that distinct PA tails of each human 19S subunit are inserted into only 5 of the 7 α-pockets due to the symmetry mismatch between the 6-fold 19S and 7-fold 20S (**Fig. 1b**). Moreover, the process of binding appears to be stepwise and dependent on the ATP hydrolysis states of the 19S. For example, the C-terminal sequence of the Rpt5 subunit: NLQYYA (**Fig. 1b**; blue square), binds first to the h20S and is essential for subsequent steps^15,17,20,22^. Following Rpt5 and Rpt3 binding to the α5-α6 and α1-α2 pockets, respectively, the other C-terminal tails bind to achieve full gate opening^22–24^. Consistent with these studies, only peptides derived from Rpt5 were found to activate h20S on their own in biochemical studies^15^. Using this platform, a recent structure-activity relationship (SAR) study identified a peptide (Ac-NLSYYT-OH) based on Rpt5 that activates the h20S proteasome by 300-fold^25^. This SAR study additionally showed that the h20S has constraints not found in archaeal or yeast proteasomes; accordingly, the C-terminal motif for the human system was termed the “YΦ motif”^25^ to differentiate it from the simpler HbYX motif of model organisms. In the YΦ motif, the C-terminal amino acid is preferred to be a small, polar amino acid (P1), followed by an interchangeable tyrosine or phenylalanine (P2) if the next amino acid is a tyrosine (P3), followed by another small, polar amino acid (P4)^25^. In this naming convention, these positions are described in reverse numerical order with the C-terminus being P1, and subsequent positions P2-P6 in the N-terminal direction.

Despite this progress in understanding the structural basis of h20S binding, it remains unclear what molecular contacts with the YΦ motif are specifically required for gate opening. Here, we show that a conserved P5 leucine is an essential part of this process. We first used hexapeptides in binding and activity assays to demonstrate that the P5 residue is critical for h20S activation, with a less pronounced impact on binding affinity. Then, to understand the structural basis for this effect, we appended a subset of these variants to PA26, a well-characterized^20,26^ protozoan PA. Specifically, the PA26 construct that we used also includes a point mutation at E102A to disable an “activation loop”, such that h20S binding and activity becomes solely dependent on the C-terminal tails^20,25^. We then solved cryo-EM structures of five PA26_E102A_ constructs bound to the h20S, showing that the P5 residue promotes gate opening by displacing a conserved R20 side chain on the h20S gate loop. Finally, we show that expression of the PA26_E102A_ fusions in a stable, inducible cell model increases total proteasome activity by approximately 2- to 6-fold, confirming the close relationship between function and gate opening. Together, these results reveal an essential role of the P5 position in h20S gate opening. These findings also show that the identity of this position can be used to “tune” total proteasome activity *in vitro* and in cells. We propose that this mechanistic knowledge will aid in the design of strategies for h20S proteasome activation.

## RESULTS

### Proteasome Activation Relies on Sequence Conserved Leucine

To identify contacts that might be essential in h20S gate opening, we first performed an evolutionary analysis of 19S subunit Rpt5. The HbYX motif is prominent in Rpt5 sequences of yeast and fungi (**Fig. 1c**; red sequence). However, in the Rpt5 tails from higher eukaryotes (**Fig. 1c**; green sequence), the distinct YΦ motif is accompanied by an invariant leucine at P5 (**Fig. 1c**; blue sequence). Previous observations showed out of all C-terminal sequences of the h19S, only the peptide derived from Rpt5 promoted 20S activity^15^. To investigate the importance of this P5 leucine, we used a standard fluorescence assay to monitor the chymotryptic activity of purified h20S. We confirmed this finding using a hexapeptide from Rpt5 (EC_50_ = 34.02 ± 15.51 μM) (**Fig. 1d**). We also confirmed that the optimized NLSYYT peptide^25^ was more a more potent activator (EC_50_ = 9.18 ± 2.25 μM). To probe binding of these peptides to the 20S, we developed a fluorescence polarization (FP) assay in which displacement is measured of a fluorescent tracer based on the NLSYYT sequence. When titrated against h20S, we calculated the apparent K_D_ of the probe to be 31 ± 1.7 nM (**Fig S1a**). When we tested the 19S-derived peptides in this competition format, we found that NLSYYT (K_I_ = 8.2 ± 2.6 μM; salmon) is a slightly better competitor than the native Rpt5 sequence (K_I_ = 13.4 ± 2.3 μM; cyan) (**Fig. 1e, S1b**). The peptides corresponding to the other 19S subunits showed no evidence of competition (K_I_ > 100 µM), consistent with the proteasome activity findings.

With these two assays for h20S activity and binding in-hand, we tested a series of peptides where the P5 position was replaced with each of the natural amino acids, plus norleucine and norvaline. Initially using a single, saturating concentration (148 μM) in the chymotryptic activity assay, we found that most P5 mutations resulted in a complete loss of activity, with the exceptions of three residues: F (61 ± 6.7%), Y (76 ± 7.0%), and W (55 ± 2.9%) (**Fig. 1f**). Even modest changes from P5 leucine to the chemically related valine, isoleucine, norleucine, or norvaline ablated activity, suggesting local, steric sensitivity. To quantify relative potency, we performed dose-response assays on the active peptides, revealing that all the EC_50_ values were increased relative to N<u>L</u>SYYT (EC_50_ = 18 ± 8.8 μM) (**Fig. 1g**). We also included the non-activating peptide, NLSYAT, as a negative control. The most active of the variants was N<u>W</u>SYYT (EC_50_ = 25 ± 3.4 μM), while the others were progressively less active. Similar potency values were obtained in assays that measure trypsin-like and caspase-like h20S activity (**Fig. S1c,d**). We also tested peptides where substitutions were introduced to the P6 position to see if this position contributes to h20S activity or binding. The results of those experiments showed minimal shifts in h20S activity, leading us to conclude that P6 is permissive whereas P5 is sensitive to changes (**Fig. S1e-g**).

We next explored the impact of the P5 position on peptide binding to h20S in an FP competition assay (**Fig S1b,h**). We found that P5 substitutions had minimal impact on binding, with K_I_ values that are comparable to leucine. For example, the K_I_ values of N<u>F</u>SYYT (15.6 ± 5.3 μM), N<u>W</u>SYYT (4.8 ± 2.1 μM), and N<u>Y</u>SYYT (11.5 ± 5.9 μM) were less than 2-fold different from N<u>L</u>SYYT (8.2 ± 2.6 μM) (**Fig. 1h, S1h**). Slightly higher K_I_ values were observed for N<u>V</u>SYYT (24.1 ± 7.15 μM) and NISYYT (43.2 ± 13 μM), indicating only minor shifts in binding. As a further control, we again tested the non-binding peptide: NLSY<u>A</u>T^25^ and confirmed no binding in this assay platform. Together, these results suggest that the P5 position is critical for h20S activation, with a less pronounced impact on binding.

### Position P5 Plays an Essential Role in Multivalent Proteasome Activation

To better understand the role of the P5 leucine in context of an intact, multivalent proteasome activator complex, we mutated the C-terminus of PA26_E102A_ to contain those peptide sequences that we had found to bind to h20S (**Fig. 2a**)^25^. The nomenclature convention of this system is based on previous work^25^, such that the fusion to the NLSYYT sequence is termed PA26_YYT_. Accordingly, we generated PA26_YYT_ variants where P5 of the C-terminal tail was replaced with phenylalanine (PA26_P5F_), tryptophan (PA26_P5W_), tyrosine (PA26_P5Y_), valine (PA26_P5V_), or isoleucine (PA26_P5I_) (**Fig. 2b**). As a further negative control, we also generated the fusion to the inactive sequence: NLSYAT (PA26_YAT_).

**Figure 2:**
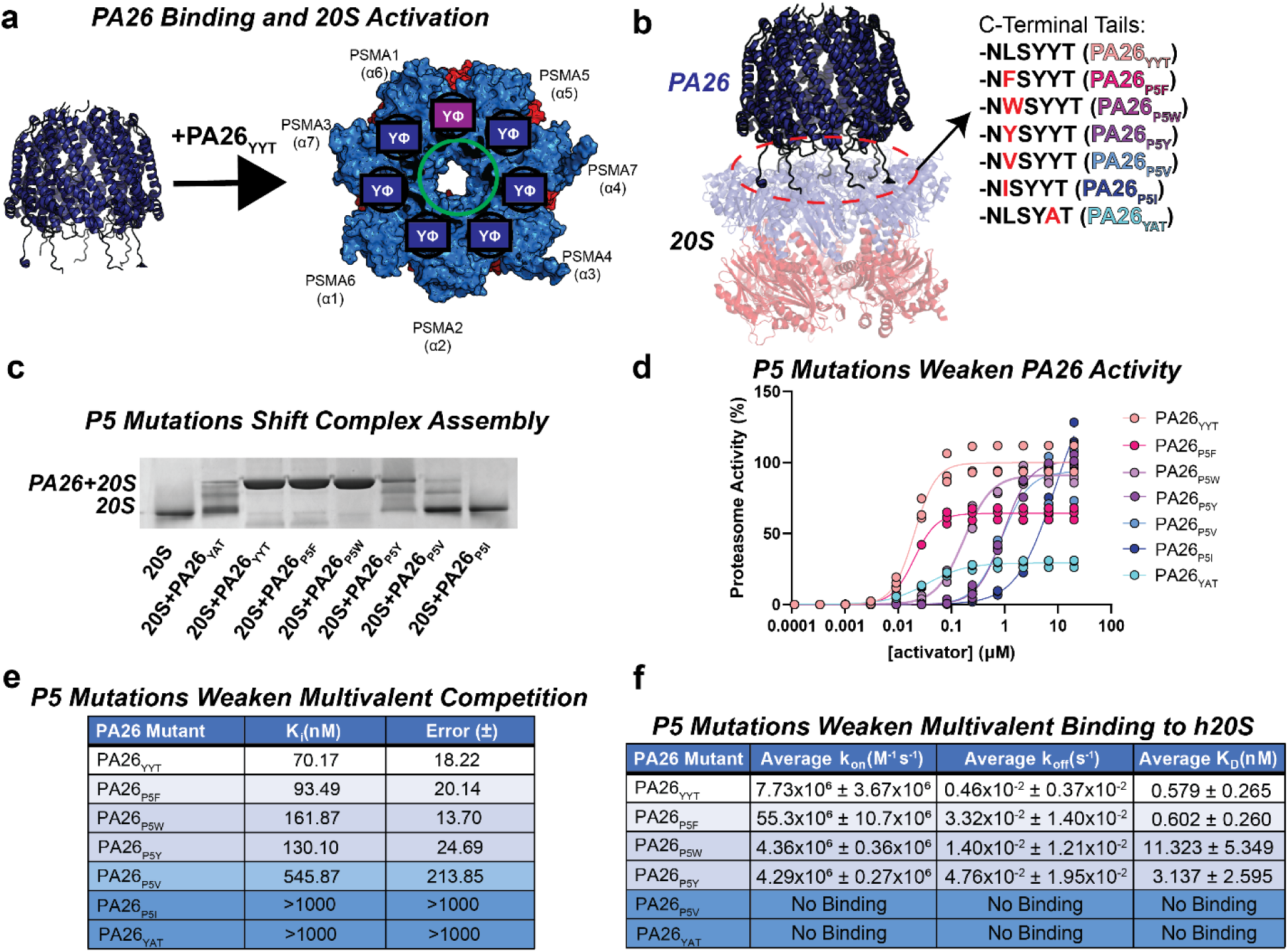
Position P5 Plays an Essential Role in Multivalent Proteasome Activation. **a)** Binding mode of PA26_YYT_ to h20S. When bound to the h20S, seven identical tails (dark blue boxes) are simultaneously inserted into all α-subunit pockets. The Rpt5 binding pocket is highlighted (purple box). **b)** Structure of PA26_YYT_ (dark blue) bound to h20S proteasome (PDBID: 6XMJ). The P5 substitution constructs were designed by replacing the native 6 C-terminal amino acids of PA26 with NLSYYT (PA26_YYT_, salmon), NFSYYT (PA26_P5F_, pink), NWSYYT (PA26_P5W_, lavender), NYSYYT (PA26_P5Y_, purple), NVSYYT (PA26_P5V_, blue), NISYYT (PA26_P5I_, dark blue), and NLSYAT (PA26_YAT_, cyan). All following figures use this color scheme. **c)** Native-PAGE of h20S proteasome incubated with each PA26 construct at 1:10 concentration ratio for 15 m before loading. **d)** Proteasome chymotrypsin-like activity assay. Maximum activity was normalized to PA26_YYT_. PA26_YYT_ (EC_50_ = 19.61 ± 5.17 nM), PA26_P5F_ (EC_50_ = 19.92 ± 2.29 nM), PA26_P5W_ (EC_50_ = 170.7 ± 17.43 nM), PA26_P5Y_ (EC_50_ = 941.6 ± 155.3 nM), PA26_P5V_ (EC_50_ = 883.4 ± 59.36 nM), and PA26_P5I_ (EC_50_ = 9142 ± 3265 nM) shown with average curve fits across individually reported data (n=3). **e)** Table of estimated K_I_ values from a fluorescence polarization competition assay. Reported values are averaged (n=3). **f)** Single concentration (1 nM PA26) surface plasmon resonance results table. Reported k_on_ and k_off_ values are averaged (n=3). Individual K_D_ values (n=3) were averaged and reported.

In the first set of experiments, we tested whether PA26 variants retained binding to the h20S. Accordingly, we purified the proteins and qualitatively studied their binding to h20S using native PAGE. Compared to a loading control of unbound h20S, we clearly visualized a band that corresponded to the PA26+20S complex at a 10-fold excess of PA26. This band was observed as the major species for PA26_YYT_, PA26_P5F_, PA26_P5Y_, and PA26_P5W_. By contrast, the PA26_YAT_ negative control, PA26_P5V_, and PA26_P5I_ showed little complex and mostly free 20S (**Fig. 2c**). This result shows that the minor K_I_ shifts observed in peptides are amplified in multivalent PAs, as expected. Together, these results suggest that binding is largely intact until the most damaging of the mutations.

We then tested each PA26 construct in turnover assays to measure their impact on the three catalytic activities of h20S. In the chymotryptic assay, we found that the most potent was PA26_YYT_ (19.6 ± 1.9 nM) followed by PA26_P5F_ (19.9 ± 1.0 nM), PA26_P5W_ (170 ± 11 nM), PA26_P5Y_ (941 ± 33 nM), PA26_P5V_ (883 ± 107 nM), and PA26_P5I_ (9014 ± 1300 nM) (**Fig. 2d**). The negative control, PA26_YAT_ (cyan), produced little total activity, as expected. The same rank order was observed in assays for the trypsin-like activity and caspase-like activity (**Fig. S2a,b**). Some of the constructs, such as PA26_P5F_, had less maximum activity (65.5%) than PA26_YYT_, for reasons that are not clear.

To quantify the impacts of these P5 modifications on binding affinity, we performed FP competition assays. When we titrated the purified PA26 mutants against a fixed concentration of h20S and FP probe, we determined that the K_I_ value of the positive control PA26_YYT_ (K_I_= 70 ± 18 nM) was within 2.5-fold of PA26_P5F_ (K_I_= 93 ± 20 nM) PA26_P5W_ (161 ± 14 nM) and PA26_P5Y_ (K_I_= 130 ± 25 nM) (**Fig. 2e, S2c**). Consistent with the native PAGE, PA26_YAT_, PA26_P5V_, and PA26_P5I_ did not bind well. Due to the lack of any assembly on the Native-PAGE and absence of measurable inhibition here, PA26_P5I_ was excluded from further experiments. To corroborate these findings in an independent platform, we then performed surface plasmon resonance (SPR) studies. Fitting the kinetic progression curves produced a similar rank order to the FP competition assays, with the strongest binder being PA26_YYT_ (K_D_= 0.58 ± 0.27 nM), followed by PA26_P5F_ (K_D_= 0.60 ± 0.26 nM), PA26_P5W_ (K_D_= 11 ± 5.35 nM) and PA26_P5Y_ (K_D_= 3.1 ± 2.6 nM). The negative control and the PA26_P5V_ and PA26_YAT_ proteins did not appreciably bind (**Fig. 2f, S2d,e**).

Together, these findings show that the P5 residue is important for proteasome activation, even in the context of an intact PA26 complex. We speculate that the modest impact of P5 on binding affinity is amplified in the context of multivalent binding, when compared to the purified peptides.

### P5 Mutations Attenuate h20S Gate Opening

To understand the molecular mechanism governing P5-dependent activation, we used cryo-EM to resolve the structures of h20S incubated with and without a subset of PA26 mutants based on our activity measurements (**Fig. 2d**). These experiments are important because the different energetic landscapes^27^ in each binding pocket on the h20S likely give rise to binding variations with seven different affinities to the same PA tail (**Fig. S2f**). Thus, only structural studies would reveal which α-pockets were bound and what contacts occurred in these pockets. All structures were aligned to an existing apo-structure of h20S^19^ (PDB: 6RGQ) and compared with the previously reported PA26_YYT_-bound fully gate-opened structure^25^ (PDB: 6XMJ). We selected optimal concentrations and stoichiometries based on our measured relative affinities (**Fig. 2f**). The resulting complexes were resolved at high resolution (2.7-3.2 Å) with one face of the h20S bound (**Fig. S3-S6**; **Supplementary Table 1**). However, for PA26_P5W_, we resolved both single-(**Fig. S4**; PDB: 12CO) and double-capped (**Fig. S4**; PDB:12CP) h20S complexes with varying strengths of PA26 density. The double-capped complex provided the strongest density and subsequent analyses of PA26_P5W_ refer specifically to this interface. We also solved the structure of h20S bound to a PA26 mutant containing the C-terminal sequence of NLSFFT (PA26_FFT_; PDB: 12CS), which we previously^25^ found binds to h20S but does not activate it (**Fig. S7**; **Supplementary Table 1**).

Through this effort, we obtained a range of structures that capture intermediate and complete open/closed states of the h20S gate along with PA tail occupancy of the various h20S activation pockets. First, we checked the activation pocket occupancy and PA tail densities as a relative indicator of stable interactions with the h20S. The number of resolved PA tail densities similarly followed activity and binding rank order with 4 of the 7 resolved in PA26_P5V_, 5/7 in PA26_P5Y_, and 6/7 in all others (**Fig 3a**). We subsequently observed that pocket occupation and the biochemical data also correlate with h20S gate opening. The least active constructs, PA26_P5V_ and PA26_P5Y_, failed to stimulate gate opening with most particles adopting a closed h20S conformation (**Fig 3a**; PDB: 12CR; PDB: 12CQ). Following the rank order of PA26 stimulatory activity, the PA26_P5W_ mutant induced partial gate opening of the h20S (**Fig 3a**; PDB: 12CP). In the next most potent stimulator complex, PA26_P5F_ induced an unambiguously open h20S conformation (**Fig 3a**; PDB: 12CN), comparable to the benchmark PA26_YYT_ structure (RMSD = 0.34) (**Fig 3a**; PDB: 6XMJ). The structure of the h20S gate remained closed when bound to PA26_FFT_, as expected because it lacks the required P2 tyrosine^25^ (**Fig S8a**). Given the progressive states of gate opening across the structures, we were able to visualize h20S gate opening using the intermediate states of the gate (**Supplementary Video 1**).

**Figure 3:**
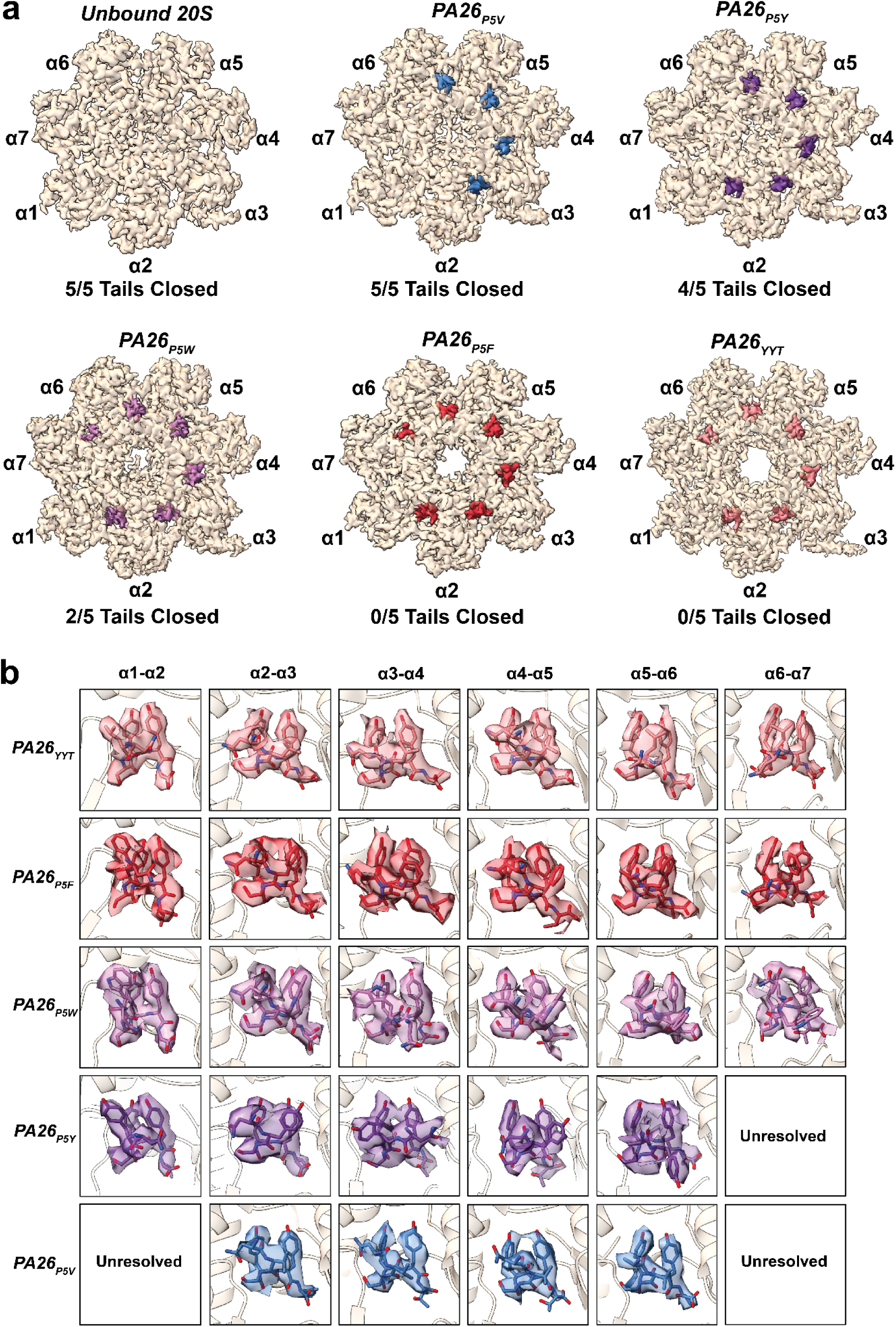
P5 Mutations Attenuate 20S Gate Opening. **a)** Top view of cryo-EM density maps of h20S either unbound (PDB: 6RGQ)^19^ or bound to PA26_P5V_ (blue, PDB: 12CR), PA26_P5Y_ (purple, PDB: 12CQ), PA26_P5W_ (lavender, PDB: 12CP), PA26_P5F_ (pink, PDB: 12CN), or PA26_YYT_ (salmon, PDB: 6XMJ)^25^. PA tails are resolved in α-subunit pockets where present. **b)** Electron density maps of PA tails shown in panel (a). Each PA tail density is shown in a grid indicating the structure and corresponding α-subunit pocket.

The number of resolved PA tails across the h20S structures follows the same rank order as the PA26 variants’ stimulatory activities, highlighting the importance of pocket occupancy. To explore this aspect further, we focused on the orientation of the PA tails in each pocket. The side chain densities of the PA tails were well resolved across all structures and enabled unambiguous modeling of the residues (**Fig. S3-S7; Supplementary Table 1**). In general, the PA tails of the P5 mutant series adopted the same tail orientations as the PA26_YYT_ benchmark in each pocket (**Fig. 3b**). By contrast, the PA tails of the PA26_FFT_ variant adopted a more relaxed conformation and adopted flexible backbone angles (**Fig S8b**). This structure highlights the importance of the parallel P2/P3 H-bonds in gate opening because the h20S gate remained closed despite occupancy of the α-pockets. However, the position of the P5 side chain was variable across structures. For example, both leucine and phenylalanine (PA26_YYT_ and PA26_P5F_) in the α3-α4 pocket orient themselves towards α4, but the residues in P5 across the other structures show side chains pointing towards α3 (**Fig. 3b**, **S8c**). These shifts might be functionally important because they correlate with the state of the h20S gate in each structure. Together, this series of five new cryo-EM structures, combined with the benchmark open and closed states, provided an opportunity to understand the molecular mechanisms underlying h20S gate opening.

### P5-Induced Shifts to the α-Subunits Retract the h20S Gate Tail

To better understand the global impact on α-subunit conformational shifts in response to the different activator tails, we measured Cα-Cα distances between the apo-structure and each PA26-bound structure. In this analysis, we considered the gate opening mechanism in a series of steps (**Fig. 4a**). Upon binding to different PA26s, the most prominent structural shift is the α-subunit ring rising, with the most distinguishable shifts around helix 1 and the gate loop^5,8^ (**Fig 4a**). Compared to the apo-structure, we see that PA26 binding variably lifts helix 1 depending on the specific α-subunit. Across the α-subunit ring, we measured the Cα-Cα distance shift of each residue across helix 1 for each structure of PA26. Overall, each PA26 universally lifts the α-subunit ring by ∼1 Å, consistent with known PA association models^23^. We can see the same activity rank order for all α-subunits except α1 and α2, where shifts are relatively equal across all PA26 mutants (**Fig. 4b**); consistent with their less critical role in gate opening.

**Figure 4:**
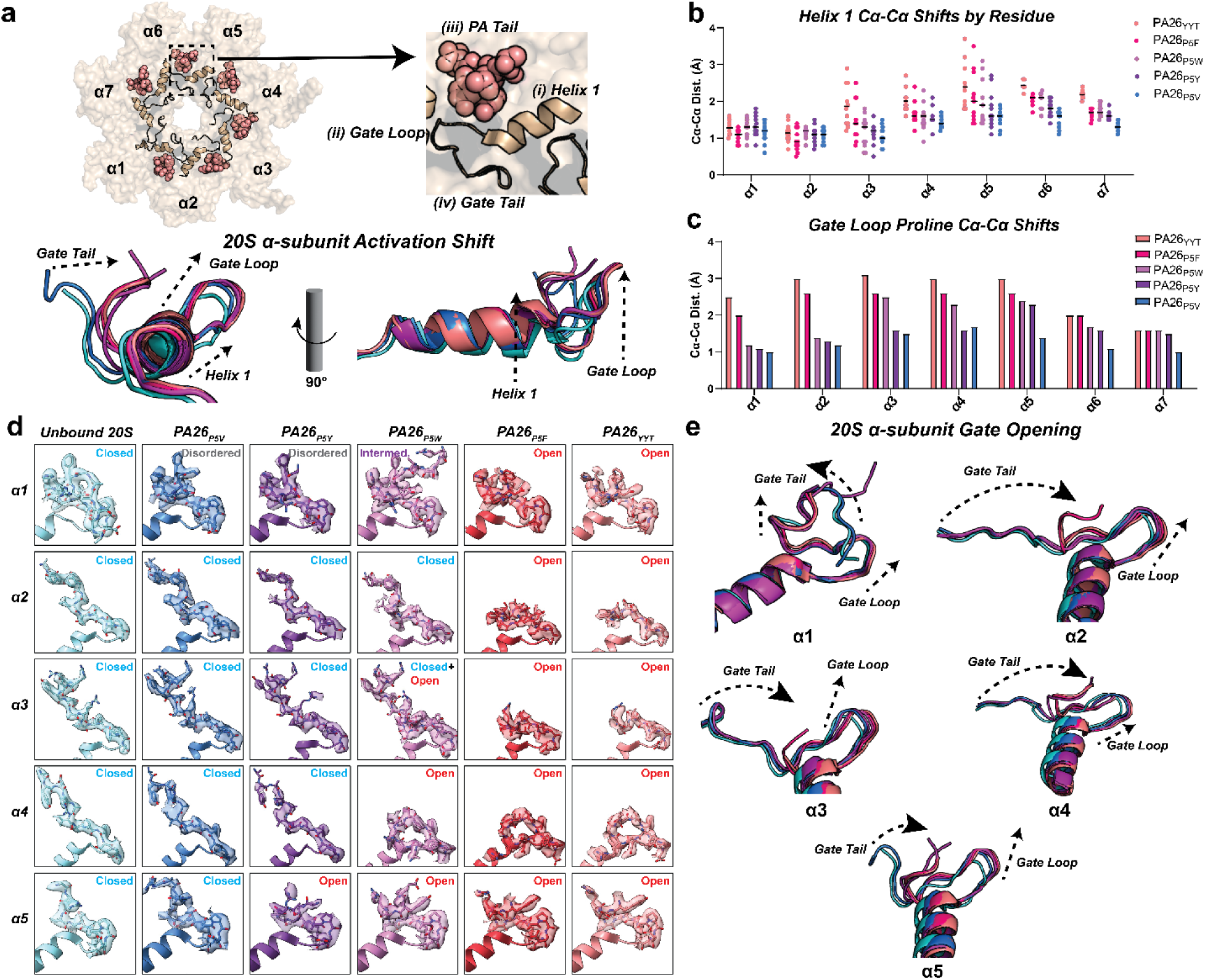
P5-Induced Shifts to the α-Subunits Retract the h20S Gate Tail. **a)** Top view of h20S bound to PA26_YYT_ (PDB: 6XMJ) with inset showing (i) Helix 1, (ii) Gate Loop, (iii) PA Tail, and (iv) Gate Tail (top). Cartoon structure of helix 1 of α5 across all structures to demonstrate observable shifts from h20S closed gate state to fully open gate state with all other PA26 bound structures (bottom). The most defining Cα shifts during gate opening can be measured by the retraction of the gate loop, and the vertical rise of helix 1. **b)** Scatter plot of Cα-Cα shifts (Å) for each amino acid in helix 1 across all structures relative to the closed-gate structure. **c)** Bar plot of Cα-Cα shifts (Å) for the conserved gate loop proline in each α-subunit. **d)** Cryo-EM density maps of 20S α-subunit gate tails across unbound h20S (light blue, PDB: 6RGQ) and h20S structures bound to PA26_P5V_ (blue, PDB: 12CR), PA26_P5Y_ (purple, PDB: 12CQ), PA26_P5W_ (lavender, PDB: 12CP), PA26_P5F_ (pink, PDB: 12CN), or PA26_YYT_ (salmon, PDB: 6XMJ)^33^. Each PA tail density is shown in a grid indicating the structure and corresponding α-subunit tail. Both observed states are shown for α3 in the PA26_P5W_-bound structure. **e)** Cartoon structure overlays of the h20S gate tails across each of the α-subunits. State shifts of the Gate Tails and Gate Loops are highlighted with arrows.

Next, we focused on PA26-induced shifts within the gate connector loops. This shift is of special interest because a conserved proline in every α-subunit is known to be a structural indicator of gate opening^4,5,8^. Accordingly, we measured the Cα-Cα distance shifts of the proline in each α-subunit across all structures (**Fig. 4c**). Out of the five gate loops, the most responsive is that of the α5 subunit. While the inactive PA26_P5V_ only displaced the gate loop by 1.4 Å, the other constructs induced shifts ranging from 2.3-3.0 Å with a prevailing rank order that again reflects their relative stimulatory activity in the biochemical assays (**Fig. 4c**). From these incremental shifts, we resolved the changes in h20S gate tail open/closed states on an atomic level (**Fig. 4d**) and relate these to the minor shifts occurring at the helix 1 interface with the PA tails. All the gate tails were unambiguously closed when bound to PA26_P5V_. In the PA26_P5Y_ structure, the h20S α5 subunit adopted an open-like state and α1 adopted a disordered state. The double capped PA26_P5W_ structure shows α5 and α4 gate tails remaining fully open and α1 being partially open in one face of the double-capped complex. Moreover, the α3 gate tail exhibited dynamic behavior with densities for both states. Finally, the PA26_P5F_ bound structure shows unambiguous gate opening across all gate tails. To open the gate tail, the gate loop extends “backwards”, while the N-terminal residues flip upwards. Although these motions are most prominent in the α5 gate tail, we see that other pockets follow this same pattern (**Fig. 4e**). Together, these observations reflect the activity data, supporting that the P5 side chain is responsible for gate opening.

### P5 Leucine Opens the Gates Through Precise Steric Displacement

To understand why P5 is important for gate opening, we focused on the α5-α6 pocket (**Fig. 5a**, left), which is associated with the initial Rpt5 interaction with the 20S^17,22,24^. We noted that in all the PA26 variant structures, the C-terminal end of the tail anchors to the α5-α6 pocket via K62 on α6, accounting for a 2 Å Cα shift on this position regardless of the open/closed state of the gate (**Fig. 5a**, right; **S9a**). Thus, we used this “anchor site” to compare the relative positions of other contacts.

**Figure 5:**
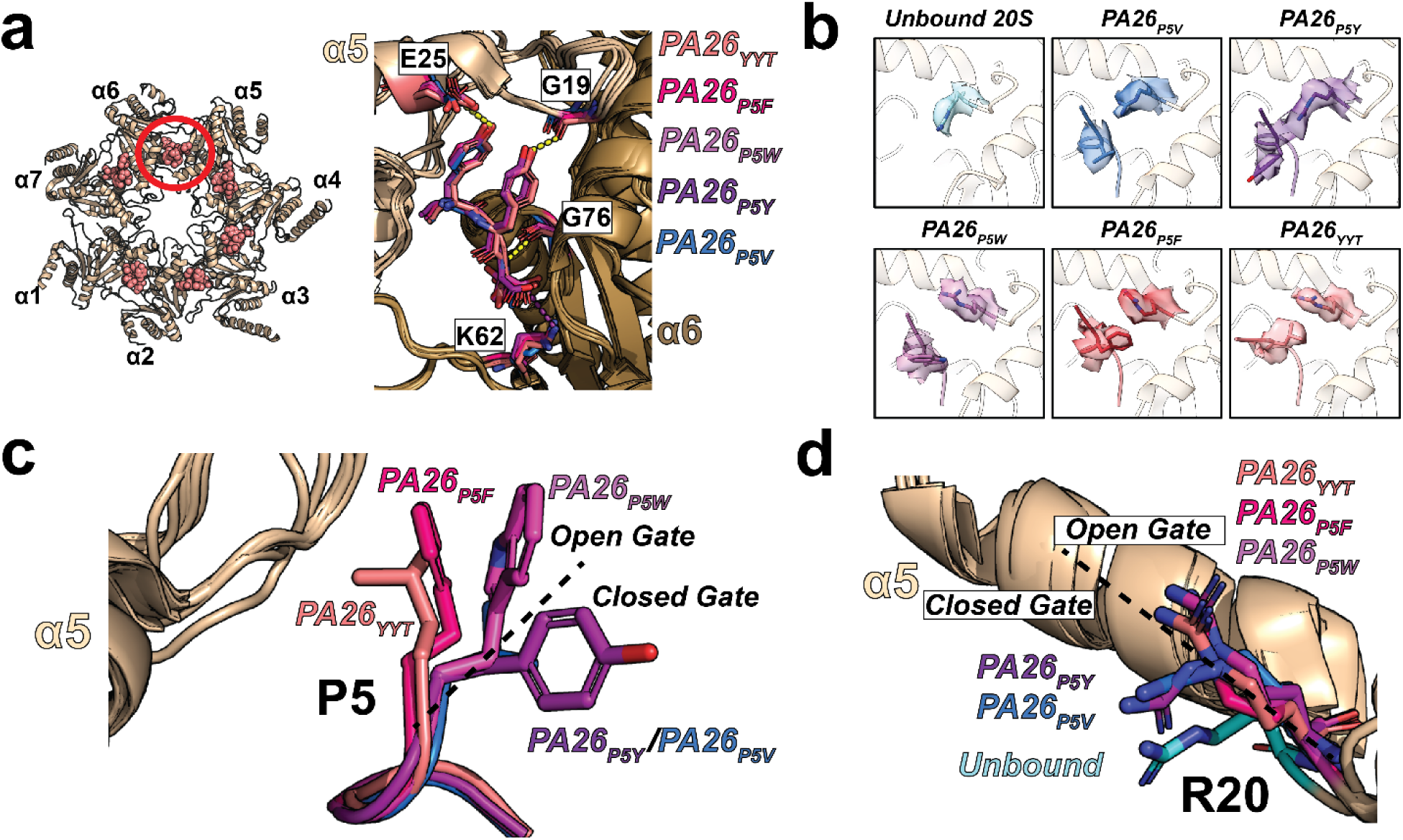
P5 Leucine Opens the Gates Through Precise Steric Displacement. **a)** Structure of the NLSYYT sequence bound within the α5-α6 pocket (left, red circle). P1-P4 side-chains of all bound PA26 constructs are shown as an overlay (right). Important H-bonds (yellow dashes) and salt bridges (purple dashes) are highlighted in the structure. **b)** Cryo-EM density maps of R20 in the 20S α5 subunit and P5 when bound to PA26_YYT_, PA26_P5F_, PA26_P5W_, PA26_P5Y_, and PA26_P5V_. **c)** P5 side-chains of each bound PA26 tail are shown as sticks alongside helix 1 of α5. All side-chain positions are shown in the majority state across particles. An arbitrary line is drawn between constructs where most gate tails are open on the left side of the line and closed on the right. **d)** Side-chains of R20 on the α5 gate loop are shown as sticks. An arbitrary line is drawn between R20 positions where most gate tails are closed vs. open.

Next, we compared the relative position of the P2 and P3 side chains. In all structures containing tyrosines at P2 and P3, we can see hydrogen bond networks bridging α5 G19 on the gate loop and α5 E25 on helix 1 (**Fig. 5a**, right). Moreover, this arrangement is stabilized by the π-π stacking between both tyrosine side chains, likely offsetting the disfavorable peptide backbone angles. This interaction was previously noted as a central component of the YΦ-motif^25^. However, these contacts are insufficient for gate opening. For example, PA26_P5V_ and PA26_P5Y_ have P2/P3 engaged, yet the gates are closed and enzymatic activity is not stimulated (**Fig. 2d**, **5a**). When all the PA26-bound structures are compared across aligned h20S particles, we find that the P2/P3 tyrosines adopt the same conformation as the PA26_YYT_ orientation (**Fig. 5a**, right). This result implies that an intact hydrogen bonding network with P2/P3 is only part of the gate opening mechanism, while variations in h20S gate opening and activity arise from elsewhere in the PA tail.

By contrast, the position of the P5 residue seems to correlate directly with gate opening. In all structures that show the h20S gate fully open (PA26_YYT_ and PA26_P5F_) or partially open (PA26_P5W_), the P5 side-chain points “inwards” towards the gate loop of α5 (**Fig. 5b**). By contrast, the P5 side-chain points “away” from the α5 gate loop in structures mostly (PA26_P5Y_) or completely closed (PA26_P5V_) (**Fig. 5b,c**). The relative position of the P5 side chain was sufficient to report h20S gate opening and activity across these structures. The most active leucine in PA26_YYT_ was oriented closest to the gate loop, followed by phenylalanine and tryptophan. These amino acids seem to open the gate through a hydrophobic/steric interaction with a single residue, R20. PA26_P5V_ and PA26_P5Y_ only lift the α5 gate loop, while the remaining PA26 constructs additionally flip the guanidinium group of R20 away from the pocket interior (**Fig. 5b,d**). This motion captures gate opening more succinctly across the α-pockets. For example, PA26_P5V_ only displaced R20 Cα by 2.0 Å while other structures shifted by 2.7-2.9 Å (**Fig. S9b**). Moreover, the displacement of the R20 guanidium group increases from a 2.0-2.6 Å shift of the R20 Cζ (guanidinium carbon) in closed structures to 4.3-4.9 Å in open structures (**Fig. S9b**).

Combined with the rigidity of helix 1, we speculate that this motion requires a downstream transfer of rotational energy within the gate loop. In other words, while the P2/P3 contacts immobilize helix 1 and G19, P5 twists the loop via R20 and subsequently pulls the gate loop “backwards”. This motion additionally results in a translational shift of P17 on the gate loop, which may contribute to allosteric energy shifts on the adjacent α6 subunit through Y24 (**Fig. S9c**), propagating the mechanism to adjacent protomers. This Pro/Tyr interface is a highly conserved feature across archaeal and eukaryotic proteasomes, and is speculated to contribute to global α-subunit ring shifts upon PA binding^4–6,8^. In the heteroheptameric rings of eukaryotic proteasomes, it is tempting to speculate that progressive shifts of the gate loop contribute to an intra-subunit allosteric network via the Pro/Tyr interface. These structures now provide us with an expanded SAR model showing how P5 tunes the h20S gate opening mechanism with the rest of the YΦ-motif.

### The P5 Position Dictates Relative Proteasome Activity in Cells

To test the putative role of the P5 position in proteasome activation in cells, we cloned each PA26 P5 mutant into pSBtet-RP vectors and transfected them into HEK293T cells to generate stable cell lines containing doxycycline-inducible PA26 variants^28^ (**Fig. 6a**). Using cell-permeable luminescent substrates, we tested for h20S catalytic activity upon dosing the cells with doxycycline (dox). We performed initial experiments to confirm that maximum proteasome activity was observed ∼48 hours after 1 μg/mL dox treatment (**Fig. S10a)** and that the expression of all PA26 constructs does not result in significant cell proliferation or toxicity effects after 48 hours (**Fig. S10b**).

**Figure 6:**
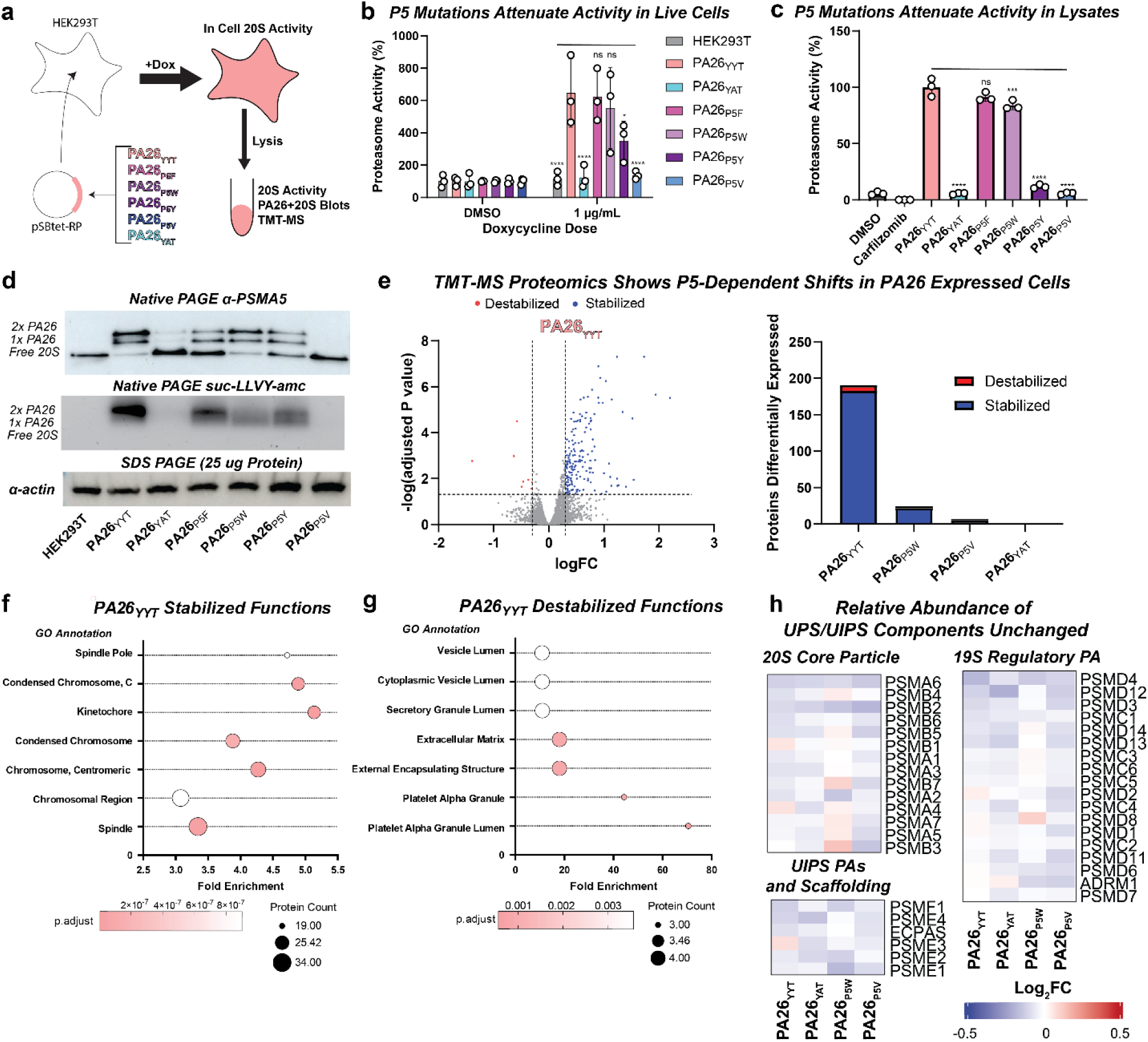
The P5 Position Dictates Relative Proteasome Activity in Cells. **a)** Schematic showing the design of cell-based experiments using the transpose sleeping beauty method^28^. **b)** Live cell proteasome chymotrypsin-like activity assay. Activity was normalized to DMSO-treated cells for all cell lines, and activity was measured for cells treated with 1 μg/mL doxycycline. Data reported individually as biological replicates (n=3). Significance values were calculated by two-way ANOVA and compared to PA26_YYT_ where ns = not significant, p<0.1 = *, p<0.01 = **, p<0.001 = ***, and p<0.0001 = ****. **c)** Proteasome chymotrypsin-like activity assay on HEK293T lysates. Data reported individually as biological replicates (n=3). Significance values were calculated by one-way ANOVA and compared to PA26_YYT_ where ns = not significant, p<0.1 = *, p<0.01 = **, p<0.001 = ***, and p<0.0001 = ****. **d)** Native PAGE western blot of 25 μg of total protein from HEK293T lysates blotted with α-PSMA5 to visualize free, single, or double PA26-capped h20S (top). An identical native-PAGE soaked in buffer containing a fluorescent activity reporter visualizes the location of active PA26+20S on the gel (middle). Loading was validated using an SDS-PAGE loading control western blot for α-actin (bottom). **e)** Volcano plot summarizing TMT-MS proteomic analyses of HEK293T lines expressing PA26_YYT_ (left). Differentially expressed hits are shown as stabilized (blue dots) or destabilized (red dots). Hits are constrained by FDR < 0.05 (horizontal dashed line) and >30% FC change (vertical dashed lines). Reported hits are consistent across biological replicates (n=3). A bar plot shows the total number of differentially expressed proteins detected across HEK293T lines expressing PA26_YYT_, PA26_P5W_, PA26_P5V_, and PA26_YAT_ (right). **f)** Enrichment of stabilized protein hits in the PA26_YYT_-expressing cell line organized by GO annotation of function. Hits were selected using the same FDR and FC constraints as (e). Number of proteins are represented by circle size, while p-value is shown by color gradient (salmon to white). **g)** Enrichment of destabilized protein hits in the PA26_YYT_-expressing cell line organized by GO annotation of function. Hits were selected using the same FDR and FC constraints as (e). Number of proteins are represented by circle size, while p-value is shown by color gradient (salmon to white). **h)** Heat maps of relative peptide abundance for proteins associated with the 20S core particle subunits, 19S regulatory PA, and UIPS PAs across the cell lines. Abundance is shown as Log_2_FC on a scale of −0.5 (blue) to 0.5 (red).

We measured proteasome function in response to PA26 expression using cell permeable luminescent probes for chymotrypsin-like, trypsin-like, and caspase-like activities. After normalizing the signal to solvent (DMSO) controls, the expression of PA26_YYT_ (646.9 ± 212.7%) showed the highest increase of total chymotrypsin-like activity, followed by PA26_P5F_ (623.5 ± 163%), PA26_P5W_ (554.5 ± 251%), and PA26_P5Y_ (351.6 ± 119.6%), while the remaining mutants PA26_P5V_ (136.6 ± 24.85%) and PA26_YAT_ (123.6 ± 70.11%) showed no significant increase compared to dox-treated parent HEK293T cells (115.5 ± 43.97%) (**Fig. 6b**). Satisfyingly, these data follow the same trend of proteasome activity observed *in vitro*. Similar results were observed with the trypsin-like (**Fig. S10c**) and caspase-like (**Fig. S10d**) assays.

To further probe the relationship between the P5 residue and proteasome activity and to provide a link to biochemical studies, we added a chymotryptic fluorescent reporter peptide to 50 μg total protein lysates from each cell line. In that assay platform, we could also treat the lysate with a proteasome inhibitor, carfilzomib, to account for signal that arises from other cellular proteases. When we normalized the resulting data to the activity of the positive control, PA26_YYT_ (100 ± 7.92%), we again observed the same rank order: PA26_P5F_ (91 ± 3.8%), PA26_P5W_ (84 ± 4.1%), PA26_P5Y_ (11 ± 1.7%), PA26_P5V_ (5.7 ± 1.0%), and PA26_YAT_ (5.6 ± 0.8%) (**Fig. 6c**). Similar results were obtained when measuring the trypsin-like (**Fig. S10e**) and caspase-like (**Fig. S10f**) activity.

As another independent test of this model, we used native-PAGE followed by blotting with an α-PSMA5 polyclonal antibody to measure the in-gel activity of the proteasome complexes^29^. As observed in our *in vitro* studies, we successfully visualized 20S proteasome capped by a single PA26 (1x) or double capped (2x) (**Fig. 6d**; top). The most active variant, PA26_YYT_ showed a bias for double capped complexes, while inactive variants PA26_YAT_ and PA26_P5V_ primarily yielded unbound h20S. The active P5 variants: PA26_P5F_, PA26_P5W_, and PA26_P5Y_, show mixtures of all 3 states. In the extreme, PA26_YYT_ seems to “soak up” the available h20S proteasome into complexes. We added the proteasome substrate, suc-LLVY-amc, to the buffer of the native-PAGE to visualize the levels of proteasome activity in each band. The resulting image confirms that the highest activity is present in the double capped band of PA26_YYT_ with less activity in the PA26_P5F_, PA26_P5W_, and PA26_P5Y_ samples and no activity in lysates containing PA26_YAT_ or PA26_P5V_ (**Fig. 6d**; center). Together, these results show that mutations to the P5 position “tune” PA26 activity in cells.

To further understand how expression of these PAs impacts the proteome, we performed TMT-MS experiments in biological triplicates using a subset of the cell lines that represent maximum, intermediate and low activation: PA26_YYT_, PA26_P5W_, PA26_P5V_, and PA26_YAT_. The total number of detected proteins across all samples ranged from 8537-8540 (**Supplementary Data 1**). We observed proteins that were either stabilized or destabilized by expression of the PA26 variants (**Fig. 6e**). Total impacted proteins followed the expected trend from the other assays: PA26_YYT_ produced the most changes; PA26_YAT_ the least. Most of these proteins are associated with chromosome stability and kinetochore organization (**Fig. 6f; Supplementary Data 2**). These proteins are related to the function of the orthologous human proteasome activator, PA28γ^30,31^, likely suggesting that over-expression of PA26_YYT_ is partially disrupting endogenous PA28γ function in this model. By contrast, the proteins that were destabilized by PA26_YYT_ expression were largely secreted and granule proteins (**Fig. 6g; Supplementary Data 2**). We speculate that excess UIPS activity might promote degradation of unfolded or unstructured polypeptide chains emerging from the secretory pathway, but it is unclear since our model uses a protozoan PA in a human background. Nevertheless, we searched for catalytic peptides to confirm h20S maturation. The N-terminal tails of the catalytic subunits β5, β1, and β2 are normally cleaved during h20S processing to expose the catalytic site (**Fig. S10g**). Thus, it is possible to search for the relative abundance of these peptides for each subunit. In this model, we found that the catalytically active peptides for β5 and β1 were unaffected by differences in PA26-enhanced activity (**Fig. S10h**). β2 peptides were not quantified. These results did not correlate with the relative abundance of factors associated with proteasome maturation or function. For example, the abundance of h20S core subunits, 19S subunits, and alternative PAs across all cell lines showed no significant differences (**Fig. 6h**). Likewise, the relative abundance of other UPS/UIPS associated proteins did not change except for three proteasome adaptor proteins (AKIRIN2, AKIRIN1, and ZFAND5) that correlated with active PA26 constructs (**Fig. S10i**). These adaptors are associated with the import of the h20S core particle into the nucleus, where the majority of UIPS PA associations have been previously observed^31,32^. This shift in protein expression would be consistent with increasing nuclear transport of h20S in response to an increased abundance of UIPS activators. Together, these proteomic studies provide strong support to a model in which the P5 position is a major determinant of proteasome activity.

## DISCUSSION

Activation of the h20S is a major step in proteostasis, allowing the regulated turnover of the proteome. While previous work has begun to unravel the molecular mechanisms of h20S gate opening in model organisms and humans, knowledge of the key contacts was incomplete. Here, we show that the highly conserved P5 leucine is an essential part of the activation model. This residue seems to be critical for contacting a conserved basic residue and opening the h20S gates, as revealed through a series of cryo-EM structures. This previously unrecognized aspect of the activation mechanism extends our knowledge, while also providing a site that can be readily “tuned” to achieve a desired level of 20S activation/activity. We find it fascinating that even a single non-enzymatic residue in a near Mega-Dalton sized protein complex, can have such a profound impact on function.

### An Advanced Molecular Mechanism for h20S Binding and Gate Opening

The compatibility of PAs with the α5-α6 pocket has been previously highlighted as one of two initial binding sites of the complete 19S PA in the UPS^16,24^, and the essential C-terminal tail of PA200^7,19^. Within this site, the essential h20S amino acids in α5 (G19 and E25) and α6 (K66) are conserved across the other α-subunit pockets. Thus, lessons learned from the α5-α6 pocket might generally translate across the h20S, a conclusion supported by our cryo-EM structures and summarized in our new SAR model (**Fig. 7**). The key distinction with our updated activation model is the inclusion of P5 side chains being important for activation of the h20S, while acknowledging that binding and activation are not fully correlated.

**Figure 7:**
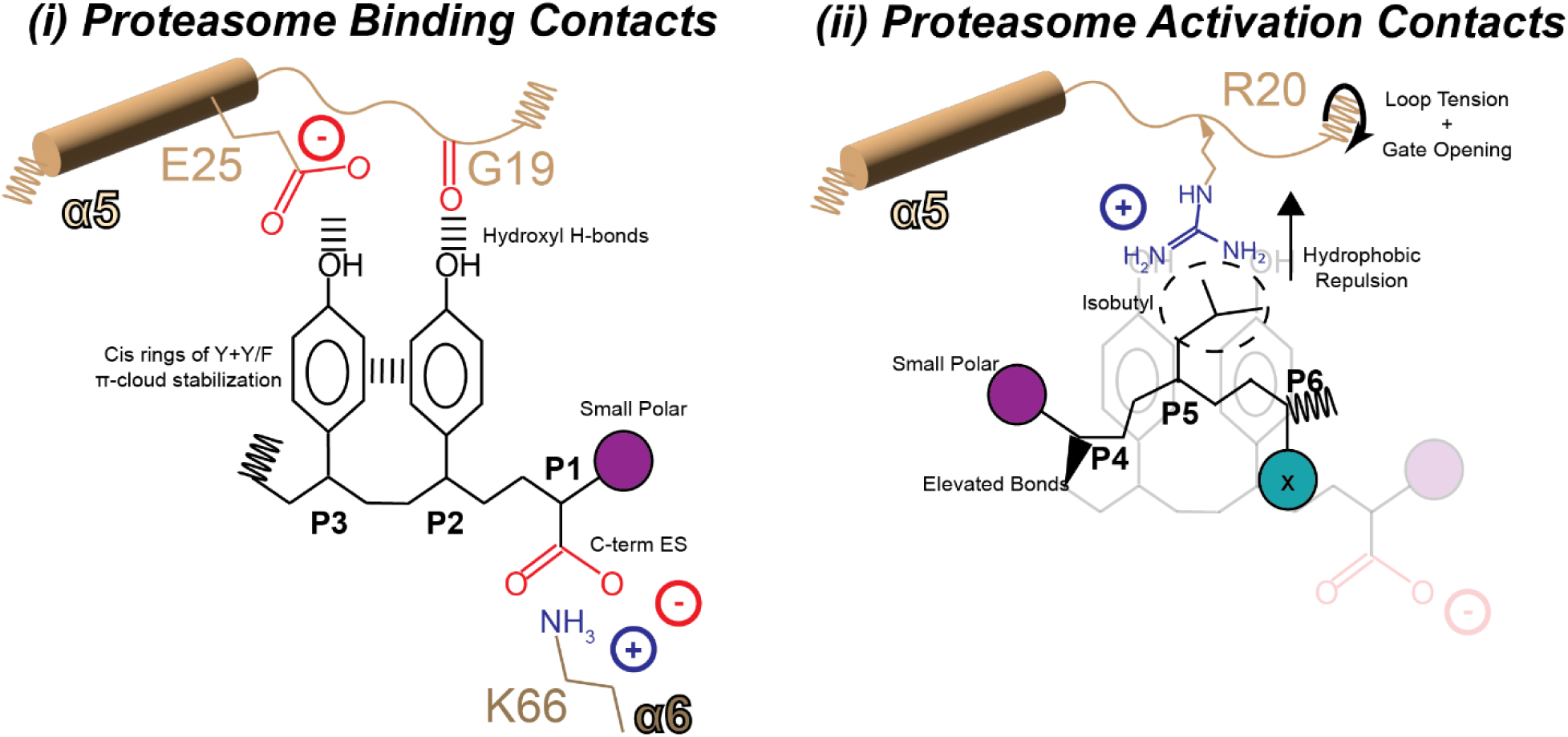
P5 Modulates Gate Opening for Substrate Selectivity. The structure-activity relationship model of optimized PA binding and h20S proteasome gate opening within the α5-α6 pocket. The model is shown in two panels to highlight interactions that are critical for PA tail binding and activation (i) and interactions that primarily contribute to activation only (ii). A free carboxyl at the P1 end forms a salt bridge with K66, with the side chain utilizing a small polar group for h20S surface compatibility. This is followed by a bulky pair of cis-aromatic side chains on P2+P3, where the π-cloud between the rings stabilizes the bulky hydrophobic groups and presents rigid hydroxyls to the α5 gate loop and helix 1 along G19 and E25. The molecule then spirals above the plane of the P2(Y/F)+P3Y so that the P5 interactions can reach R20. A small polar group in P4 assists with h20S surface compatibility and directs the sequence to spiral upwards. P5L completes the activation mechanism through its charge-neutral hydrophobic repulsion of the R20 side chain, providing the additional energy along the backbone to complete gate tail retraction while providing stabilizing hydrophobic cover over the P3Y+P2(Y/F) component below.

Our revised model reinforces two conclusions that were made previously about binding h20S: (i) the carboxyl group anchoring the protein to K66 within the pocket, and (ii) the presence of two aromatics within the YΦ-motif. While those interactions seem critical to binding, they do not define gate opening as explicitly shown in the PA26_FFT_ structure. The ability to stimulate gate opening arises from a series of interactions, including H-bonding at the P2 and/or P3 positions (**Fig. 7**). These rigidified hydrogen bond donors are presented to the G19 and E25 acceptors along the α5 gate loop and helix 1, as seen in our structures. Our results show that the leucine side chain at P5 is essential to complete the activation mechanism, while substitutions to P5 largely abolish activity. The key interaction is contact with R20, which flips the side chain backwards (**Fig. 7**). These mechanistic observations are also consistent with evolution, where R20 was present in archaeal homoheptameric 20S proteasomes^21^ before eukaryotic divergence to heteroheptameric 20S proteasomes.

Our series of cryo-EM studies with experimental characterization allows us to ask how many α-pockets must be occupied and how much gate opening is enough for substrates? We observed a wide range of relative α-pocket occupancies and can correlate them with the extent of gate opening and enzymatic turnover data. One surprising finding from this analysis is that some structures show relatively minimal gate opening, yet relatively potent stimulation. For example, PA26_P5W_ yielded single- and double-capped complex structures with varying degrees of gate opening despite having observable activity both *in vitro* and in cells. Yet, activity generally increased with the extent of gate opening, affirming that occupancy of at least 3 pockets and some gate motions are required. It is further tempting to consider what implications gate stability and minimal gate activation may have on substrate entry, where UIPS-like gate opening can be controlled to select for more disordered proteins over those with more structural elements.

### Implications for the Pursuit of Therapeutics

UIPS complexes become more abundant with age and oxidative stress^3,5,32^. Because UIPS complexes do not have unfoldase activity, they have been proposed to be especially important in the clearance of IDPs. Thus, elevated UIPS activity might be a protective response to the accumulation of IDPs in aging and disease^9,33,34^. Efforts have been previously made to direct some disordered proteins to the proteasome for degradation, such as PROTACs, but these leverage the components, such as ubiquitin ligases, within the UPS instead of the UIPS^35,36^. This approach could be problematic because UPS activity, unlike the UIPS, declines with age. If the UIPS is more compatible with IDP clearance and removal of oxidative damaged proteins^10^, then the foundation of a degradation strategy designed to clear these proteins might be better served by being centered around h20S gate opening. Pioneering work has pursued small molecule activators of the h20S proteasome^11,25,37^. The improved SAR model expands our fundamental understanding of the mechanisms of h20S activation, drawing special attention to the need for a P5-residue equivalent in any activation strategy that takes advantage of the natural stimulatory pathway.

## ACKNOWLEDGEMENTS

This work was supported by grants from the Alzheimer’s Association Research Fellowship AARF-23-1150444 (to B.D.R), Hevolution Foundation (to J.E.G. and M.A.P.) and the Tau Consortium (to J.E.G and J.S). The ALBORADA Drug Discovery Institute is core funded by Alzheimer’s Research UK (registered charity no. 1077089 and SC042474) under ARUK-2021DDI-CAM, with support from the ALBORADA Trust. Project “CP21/00017”, funded by Instituto de Salud Carlos III (ISCIII) and co-funded by the European Union (M.A.P.); Project “PID2022-136403OA-I00” funded by MCIN/AEI/10.13039/501100011033/FEDER, UE (M.A.P). TMT-MS data were acquired using instruments from the proteomics core facility at the ISPA Institute. The authors thank the laboratory of P. Coffino (The Rockefeller University) for the original PA26 construct. We also thank J. Day (UCSF) and H. Williams (ADDI) for helpful suggestions and discussions on the manuscript.

## AUTHOR CONTRIBUTIONS

B.D.R and J.E.G designed the studies and wrote the manuscript. All authors edited the manuscript. B.D.R conducted all PA26 purification, proteasome activity assays, fluorescence polarization assays, OpenSPR experiments, cell assays, and gels/blots. B.D.R generated the plasmids for dox-inducible PA26 expressing HEK293T cell lines and the cell lines themselves. A.L, H.F, and S.A designed and synthesized the FP probe used in fluorescence polarization experiments. N.L.Y and B.D.R performed cryo-EM sample preparation. N.L.Y performed cryo-EM data collections and data processing. J.R.B performed cryo-EM sample preparation, data collections and data processing on the PA26_FFT_ structure. Cryo-EM equipment and data analysis servers are managed by A.A.M and E.T. D.T-V ran and processed the TMT-MS samples. P.M-B ran bioinformatic analyses on the TMT-MS results. M.A.P aided with the interpretation of the TMT-MS results. J.E.G, B.D.R, J.S, D.R.S, and M.A.P provided funding

## DATA AVAILABILITY

The structural data generated in this study have been deposited into the Protein Data Bank (PDB) and Electron Microscopy Data Bank (EMDB). The accession codes are h20S•PA26_P5F_: PDB 12CN, EMDB EMD-76309; h20S•PA26_P5W_: PDB 12CO, EMDB EMD-76310; h20S•(PA26_P5W_)_2_: PDB 12CP, EMDB EMD-76311; h20S•PA26_P5Y_: PDB 12CQ, EMDB EMD-76312; h20S•PA26_P5V_: PDB 12CR, EMDB EMD-76313; h20S•PA26_FFT_: PDB 12CS, EMDB EMD-76314. Publicly available data used in this study include: cryo-EM structure of human 20S proteasome (PDBID: 6RGQ) and cryo-EM structure of h20S bound to PA26_YYT_ (PDBID:6XMJ). Quantitative TMT mass spectrometry data are available in Supplementary Data 1. GO annotation assignments and data are available in Supplementary Data 2. The code used for the analysis of the proteomic data is available at https://github.com/prado-lab/collab-gestwickilab-pa26-2026. All mass spectrometry proteomic data have been deposited in the ProteomeXchange Consortium via the PRIDE repository^38^ under the identifier PDXXXX. The raw and processed data generated by this study are provided in the Source Data file. All data supporting the findings of this study are available from the corresponding author upon reasonable request. Source data are provided with this paper.

## METHODS

### Reagents

Human 20S proteasome used for *in vitro* experiments was purchased from R&D Systems.

### Strains and Plasmids

The *E. coli* strain XL1-blue was used for propagating plasmids. Rosetta(DE3)pLysS cells were used for the expression and purification of all PA26_E102A_ constructs cloned into pMCSG7 vectors. pSBtet-RP^28^ was a gift from Eric Kowarz (Addgene plasmid #60497; http://n2t.net/addgene:60497; RRID:Addgene_60497). P5 mutant constructs in pMCSG7 were generated using site directed mutagenesis. SB100X^39^ in pCAG globin pA was a gift from Mark Groudine (Addgene plasmid # 127909; http://n2t.net/addgene:127909; RRID: Addgene_127909). Doxycycline-inducible PA26 expression cell lines were generated by cloning the genes of each PA26 gene into the pSBtet-RP vector, which was used during transfection along with a helper plasmid pSB100X. Primers used to generate PA26 P5 mutants are listed in Supplementary Table 2. Primers used to clone PA26 constructs into pSBtet-RP vector are listed in Supplementary Table 3.

### Peptide Synthesis

All peptides were synthesized and purchased from Genscript USA. Complete sequences and terminal modifications are listed in Supplementary Table 4.

### Proteasome Fluorescent Activity Assay

Human 20S proteasome (R&D Systems, E-360) activity was measured in 384-well plates (Griener Bio-One, 781209) using fluorgenic substrates suc-LLVY-amc (AnaSpec, AS-63892), boc-LRR-amc (AdipoGen, AG-CP3-0014), and FAM-LFP (5-FAM-AKVYPYPMEK(QXL520)-NH_2_; AnaSpec) to measure chymotrypsin-like, trypsin-like, and caspase-like activities, respectively. All experiments were performed in assay buffer (50 mM Tris pH 7.5 1 mM DTT, 0.01% Pluronic F-68 (Gibco Life Technologies, 24040032) in volume of 30 μL per sample across technical triplicates. Human 20S (final conc. 4 nM) was incubated in the presence or absence of activators (Final conc. Peptides: 0.2-148 μM; PA26: 1.1x10^-4^-20 μM) at room temperature for 5 min. Substrate (10 μM suc-LLVY-amc; 20 μM boc-LRR-amc; or 100 nM FAM-LFP) was added to each sample immediately before reading. Fluorescence intensity was measured (suc-LLVY-amc and boc-LRR-amc Ex: 355 nmn Em: 440 nm; or FAM-LFP Ex: 490 nmn Em: 520 nm) on a Spectramax M5 microplate reader (Molecular Devices; SoftMax Pro 6.5.1). Values measured were in relative fluorescence units (RFU). The hydrolysis rates were calculated from the slopes of data between 100-500 s (suc-LLVY-amc and FAM-LFP) or 500-1000 s (boc-LRR-amc).

All data analysis and statistics were calculated and plotted using GraphPad Prism 10.6.1. Baseline hydrolysis was normalized to the total mean activity for the lowest concentration of every activator assayed in each plate and the maximal hydrolysis was normalized as reported. Normalized activity was plotted relative to log_10_(activator) and fit to the model for log(agonist) vs. response (variable slope). Final activity values for each concentration of activator were calculated by inserting the values of the previous fit into Eq. (1), where X= log[activator].

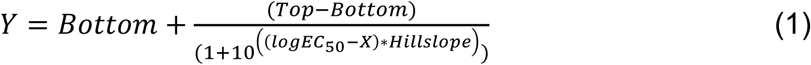

The final X, Y values were plotted and the curve fitted to the model for Sigmoidal, 4PL. The estimated EC_50_ of each activator was calculated based on these curve fits.

### Fluorescence Polarization

The fluorescence polarization probe (FP probe) used in this study is the property of the ALBORADA Drug Discovery Institute, and specific details regarding the design and synthesis of the probe are not discussed here. Fluorescence polarization results were measured in 384-well plates (Griener Bio-One, 781209). The initial binding experiment between h20S and the FP probe was assembled in assay buffer (50 mM Tris pH 7.5 1 mM DTT, 0.01% Pluronic F-68 (Gibco Life Technologies, 24040032), where a final concentration of 0.4 nM FP probe was added to a concentration gradient of 120-0.02 nM h20S. Each h20S sample was run as technical triplicates. All samples were incubated for 15 m at room temperature before taking readings. The samples were read using a BioTek Synergy H1 plate reader (BioTek) equipped with a FITC filter cube (Ex: 498 nm Em: 517 nm). The data was plotted and analyzed in GraphPad Prism 10.6.1 and fitted to the [Inhibitor] vs. response (three parameters) model.

Fluorescence polarization competition assays were performed using the same equipment and reagents as above but including activators. A fixed concentration of 8 nM h20S was added to a gradient of activators (peptides: 50-0.024 μM; PA26: 320-0.156 nM) and incubated for 15 m at room temperature. Each sample was run as technical triplicates. Before readings, 0.4 nM FP probe was added to all samples and mixed.

### PA26 Purification and Expression

All PA26 constructs were generated in pMCSG7 vectors and transformed into *E. coli* Rosetta2(DE3)pLysS cells for recombinant expression and purification. PA26_E102A_ constructs PA26_YYT_ and PA26_YAT_ were generated as previously described^25^. P5 mutant constructs PA26_P5F_, PA26_P5W_, PA26_P5Y_, PA26_P5V_, and PA26_P5I_ were generated using site-directed mutagenesis using the PA26_YYT_ construct as a template. *E*. *coli* cells were grown overnight in 20 mL terrific broth (TB) with 1x ampicillin at 37°C. In the morning, the overnight culture was added to 1 mL TB with 1x ampicillin and incubated at 37°C, induced with 1 mM isopropyl β-d-1-thiogalactopyranoisde (IPTG) in log phase, cooled to 18°C, and grown overnight. Cells were harvested by centrifugation at 5k xg for 15 min, resuspended in His resin binding buffer (20 mM Tris pH 7.9, 20 mM NaCl, 5 mM imidazole, supplemented with EDTA-free cOmplete™ protease inhibitor cocktail (Roche)), sonicated (35% power; 30 s pulse; 1 min rest) for 10 min, and clarified by centrifugation at 45k xg for 20 min. Clarified lysate was applied to Ni-NTA His-Bind Resin (Novagen). The resin was washed in 35 mL wash buffer (20 mM Tris pH 7.9, 20 mM NaCl, 5 mM imidazole) and eluted in a gradient of elution buffer (50-200 mM imidazole). Purified protein was treated with 1 mM DTT for 1 hr at ambient temperature and imidazole was removed by overnight dialysis into storage buffer (20 mM Tris pH 8.0, 200 mM NaCl).

### Native-PAGE Activity Assays

Native PAGE was performed using NuPAGE™ 3-8% Tris-Acetate gels (Invitrogen) to optimally visualize the 20S proteasome as previously described^29^. For HEK293T cell lysates, 25 μg total protein was added in each lane. The samples were loaded onto the gel and run at 150 V for 4 h to align the h20S across the center of the gel. For in gel-activity assays, the gel was placed in a dark box containing 25 mL of reaction buffer (50 mM Tris pH 7.5, 10 mM MgCl_2_, 1 mM ATP, 1 mM DTT, 48 μM suc-LLVY-amc) and incubated at 37°C for 30 m. The gel was subsequently imaged for amc fluorescence (Ex: 355 nmn Em: 440 nm). Otherwise, gels were stained with Coomassie for protein visualization and imaged.

### Western Blotting of Native-PAGE and SDS-PAGE Gels

All primary antibodies used in western blots are listed in Supplemental Table 5. For HEK293T cell lysates, 25 μg total protein was added in each lane. For recombinant samples, 20 μg of h20S was mixed with or without 10 μM PA26 construct and incubated at room temperature for 15 m. Upon completion of running either Native-PAGE or SDS-PAGE gels, selected gels for western blotting were transferred onto a 0.2 μm mini nitrocellulose membrane (Bio-Rad) and loaded into a Trans-Blot® Turbo™ Transfer System (Bio-Rad) and run using the High MW transfer protocol for semi-dry transfer. The blots were removed and incubated on an orbital shaker in 10% non-fat milk for 1 h at room temperature. Primary antibodies were added at a 1:2000 ratio and the blots were soaked in primary antibody solution overnight at 4°C. The blots were washed with 1xTBST three times at 15 m per wash and soaked in secondary antibody (α-rabbit IgG-HRP, polyclonal goat, Jackson ImmunoResearch #111-035-144) solution at a ratio of 1:5000 for 45 m at room temperature. The blots were washed with 1xTBST three times at 5 m per wash and once with 1xTBS for 5 m. 1 mL of Amersham™ ECL reagent (Cytiva) was soaked onto the blots and the blots were subsequently imaged.

### Single Concentration Kinetics Open Surface Plasmon Resonance

All experiments were performed using a 2-channel OpenSPR^®^ instrument purchased from Nicoya Lifesciences. 1 μM h20S was biotinylated using EZ-Link™ NHS-Biotin (Thermo Scientific) and immobilized to a high sensitivity biotin-streptavidin OpenSPR sensor chip (Nicoya Lifesciences) using the manufacturer’s protocol for chip activation and ligand immobilization. Immobilization was performed using binding buffer (20 mM HEPES pH 7.5, 150 mM NaCl, 1 mM DTT, 0.01% Pluronic F-68) and blocked using binding buffer supplemented with 1% BSA (Sigma-Aldrich). PA26 constructs were diluted in binding buffer to concentration of 1 nM and injected onto the OpenSPR instrument as technical triplicates for each concentration. Replicate averaging, association and dissociation curve fitting, rate constants estimation, and K_d_ estimation were performed GraphPad Prism 10.6.1.

### Cryo-Electron Microscopy Sample Preparation and Data Collection

All h20S•PA26 complexes except for h20S•PA26_FFT_ were prepared by incubating 400 nM h20S proteasome with either 2- or 10-fold molar ratios of PA26 for 1 hour at 37 °C in vitrification buffer (50 mM Tris pH 7.5, 1 mM DTT, 0.01% Pluronic F-68). For PA26_P5F_, a 2-fold ratio was used, while remaining constructs used a 10-fold ratio. Afterwards, 3 µL of sample was applied to a non-glow-discharged graphene oxide (GO)-coated holey carbon grid (Quantifoil R1.2/1.3 on a gold 300 mesh support), which were prepared in house as previously described^40^. After 30 s, grids were blotted for 4 s at 4 °C and 100% humidity using a FEI Vitrobot Mark IV (Thermo Fisher Scientific), followed by plunge freezing in liquid ethane. For h20S•PA26_FFT_, the complex was prepared by incubating 2 µM h20S with 4 µM PA26_FFT_ in buffer (50 mM Tris pH 7.5, 10 mM MgCl_2_, 1 mM DTT), Afterwards 3 µL of sample was applied to a glow discharged (PELCO easiGlow, 15 mA, 2 min) holey carbon grid (Quantifoil R1.2/1.3 on a gold 300 mesh support), and plunge frozen in liquid ethane using the same conditions as the other samples.

For each sample except for h20S•PA26_FFT_, movies were collected at a nominal magnification of 105,000× (physical pixel size: 0.417 Å/pixel) on a Titan Krios (Thermo Fisher Scientific, Waltham, MA) operated at 300 kV and equipped with a K3 direct electron detector and BioQuantum energy filter (Gatan, Inc., Pleasanton, CA) set to a slit width of 20 eV. A defocus range of −0.8 to −1.8 μm was used with a total exposure time of 2.024 s fractionated into 0.025-s subframes. The total dose for each movie was 46 electrons/Å^2^. Movies were motion corrected using MotionCor2^41^ in Scipion^42^ and were Fourier cropped by a factor of 2 to a final pixel size of 0.834 Å/pixel. For h20S•PA26_FFT_, movies were collected in super-resolution mode at a calibrated magnification of 53,937x (physical pixel size: 0.927 Å/pixel) on a Glacios TEM (Thermo Fisher Scientific) operated at 200 kV and equipped with a K2 Summit direct electron detector (Gatan). A defocus range of −0.5 to − 2.5 μm was used for a total exposure time of 10 s fractionated into 0.1-s subframes. The total dose of each movie was 66 electrons/Å^2^. Movies were motion corrected using MotionCor2^41^ in Scipion^42^ and were Fourier cropped by a factor of 2.

### Cryo-EM Data Processing

Data processing schematics for each sample are presented in Figures S3-S7. Statistics are reported in Supplementary Table 1. For each dataset, image processing was performed in CryoSPARC v4^43^. The h20S•PA26_P5Y_ dataset additionally had some processing performed in RELION v4^44^.

For the h20S•PA26_P5F_ dataset, 6,361 motion-corrected and dose-weighted micrographs were subject to Patch CTF estimation and curated to low-resolution micrographs (CTF fit resolution > 4 Å), or those with significant ice contamination, resulting in 5,612 remaining micrographs. Particles were picked using Blob Picker and extracted with a box size of 600 pixels binned to 100 pixels for an initial particle count of 1,805,503. Two rounds of reference-free 2D classification were performed to remove contaminants and low-resolution classes, resulting in 300,401 remaining particles. These were subject to *ab initio* model generation (K=5) and heterogenous refinement using the *ab initio* models as references. Two class (97,311 particles total) displayed density corresponding to a bound PA26, while the others were not bound. Particles in the bound classes were pooled and re-extracted with a box size of 600 pixels binned to 400 pixels. The particle stack then underwent homogenous refinement followed by non-uniform refinement with CTF and defocus refinement without imposing symmetry to give the final h20S•PA26_P5F_ map at an estimated global resolution of 2.57 Å.

For the h20S•PA26_P5W_ dataset, 11,224 motion-corrected and dose-weighted micrographs were subject to Patch CTF estimation and curated to low-resolution micrographs (CTF fit resolution > 4 Å), or those with significant ice contamination, resulting in 10,031 remaining micrographs. Particles were picked using Template Picker (using h20S•_P5V_ averages as templates) and extracted with a box size of 600 pixels binned to 100 pixels for an initial particle count of 817,448. Two rounds of reference-free 2D classification were performed to remove contaminants and low-resolution classes, resulting in 336,680 remaining particles. These were subject to *ab initio* model generation (K=5) and heterogenous refinement using the *ab initio* models as references. One class (83,523 particles) displayed density corresponding to a single bound PA26, while another (81,539 particles) had two bound PA26. This single-bound particle stack was re-extracted with a box size of 600 pixels binned to 400 pixels. The single-bound particle stack underwent homogenous refinement followed by non-uniform refinement with CTF and defocus refinement without imposing symmetry to give the final h20S•PA26_P5W_ map at an estimated global resolution of 2.53 Å. We noted that the PA26 density was incomplete but did not pursue further data processing. The double-bound particle stack underwent an identical refinement workflow give the final h20S•(PA26_P5W_)_2_ map at an estimated global resolution of 2.57 Å. We noted that one face of the 20S exhibited stronger PA26 density than the other face.

For the h20S•_P5Y_ dataset, 13,698 motion-corrected and dose-weighted micrographs were subject to Patch CTF estimation and curated to low-resolution micrographs (CTF fit resolution > 4 Å), or those with significant ice contamination, resulting in 10,679 remaining micrographs. Particles were picked using Template Picker (using h20S•_P5V_ averages as templates) and extracted with a box size of 600 pixels binned to 100 pixels for an initial particle count of 411,279. Two rounds of reference-free 2D classification were performed to remove contaminants and low-resolution classes, resulting in 186,686 remaining particles. These were subject to *ab initio* model generation (K=3) and heterogenous refinement using the *ab initio* models as references. After four rounds of heterogenous refinement, one class (128,439 particles) displayed moderate density corresponding to the bound PA26. This particle stack was re-extracted with a box size of 600 pixels binned to 400 pixels. It then underwent homogenous refinement followed by non-uniform refinement with CTF and defocus refinement without imposing symmetry to give a h20S•PA26_P5V_ map with weak PA26 density. This particle stack was then exported into RELION and a mask around the PA26 was constructed. One round of 3D classification without image alignment (K=10, T=40) was performed. One class (21,162 particles) exhibited clear PA26 density. Particles in this stack were imported back to CryoSPARC and underwent non-uniform refinement with CTF and defocus refinement without imposing symmetry to give the final h20S•PA26_P5Y_ map at an estimated global resolution of 3.03 Å.

For the h20S•PA26_P5V_ dataset, 8,691 motion-corrected and dose-weighted micrographs were subject to Patch CTF estimation and curated to low-resolution micrographs (CTF fit resolution > 4 Å), or those with significant ice contamination, resulting in 8,209 remaining micrographs. Particles were picked using Blob Picker and extracted with a box size of 600 pixels binned to 100 pixels for an initial particle count of 405,101. Two rounds of reference-free 2D classification were performed to remove contaminants and low-resolution classes, resulting in 100,667 remaining particles. These were subject to *ab initio* model generation (K=3) and heterogenous refinement using the *ab initio* models as references. One class (78,669 particles) displayed density corresponding to the bound PA26. This particle stack underwent another round of heterogenous refinement (k=3) to remove unbound 20S particles, resulting in 24,078 remaining particles. This particle stack was re-extracted with a box size of 600 pixels binned to 400 pixels. It then underwent homogenous refinement followed by non-uniform refinement with CTF and defocus refinement without imposing symmetry to give the final h20S•PA26_P5V_ map at an estimated global resolution of 2.78 Å.

For the h20S•PA26_FFT_ dataset, 3,949 motion-corrected and dose-weighted micrographs were subject to Patch CTF estimation. Particles were picked using Template Picker and extracted with a box size of 600 pixels binned to 100 pixels. 2D classification, *ab initio* modeling, and 3D classification were performed, which resulted in three classes with density attributable to PA26 in addition to the 20S core particle, while two classes only had 20S density. Particles from the three PA26-bound classes were re-extracted with a box size of 600 pixels binned to 400 pixels, and jointly refined using non-uniform refinement, with local and global CTF refinement enabled. This resulted in the final h20S•PA26_FFT_ map at an estimated global resolution of 3.05 Å.

### Model building

Initial model building was performed by fitting halves of existing apo h20S (PDB: 6RGQ) or PA26_YYT_-bound h20S (PDB: 6XMJ) models into the refined h20S•PA26_P5X_ maps. Subsequent model fitting and refinement was performed using COOT^45^ and ISOLDE^46^. After Ramachandran parameters, rotamers, and clashes were satisfied, the ISOLDE command “isolde write phenixRsrInput # <model><map resolution> # <map>” was used to export the model and generate a rigid-body refinement settings file for real-space refinement in PHENIX^47^, giving the final refined models.

### Generating Dox-Inducible PA26 Expression Cell Lines

PA26-expressing cell lines were generated using a modified protocol from Kowarz, E. et al^28^. HEK293T cells at third passage cycle were plated in 10 cm nucleon delta surface dishes (Thermo Scientific) and grown in complete media (CM) containing Dulbecco’s Modified Eagle Media (DMEM; Gibco 11965-092) with 10% FBS (Gibco A5670701) and 0.01x Pen-Strep (Gibco 15140-122) to 75% confluency. In tubes for each construct, 50 ng pSB100X transposase helper plasmid and 950 ng pSBtet-RP vector containing a PA26 construct gene were mixed with Lipofectamine 3000 and P3000 reagent (Invitrogen L3000001) in Opti-MEM (Gibco 31985062) following the manufacturer’s protocol for transfection. HEK293T cells were passaged once again, and 2x10^5^ cells were plated into 100 μL CM per well in a 24-well plate (Costar 3337). 100 μL of the DNA+lipofectamine mixture was added into 3 biological replicates prepared for each PA26 construct and the cells were allowed to incubate for 24 h at 37°C. On the following day, the media was exchanged for 100 μL fresh CM containing 1 μg/mL puromycin (Gibco A1113803) and allowed to incubate for 3 d at 37°C to select for cells containing the integrated transposon with doxycycline-inducible PA26 and constitutively expressed PuroR. Upon completion of selection, the cells were passaged into fresh CM within a 10 cm plate and cultivated with standard passaging for 2 weeks to recover the cells before use in experiments.

### Mammalian Cell Lysis

For experiments using cell lysates, PA26 cell lines in 10 cm plates were treated with 1 μg/mL doxycycline and incubated for 48 h so that the cells would be harvested at 80% confluency. The cells were washed twice with ice cold 1xPBS before being scraped into 10 mL of 1xPBS and transferred into a 15 mL tube. The cells were spun at 1,500 xg for 5 m and the PBS was removed before resuspending the cells in 1 mL of 1xPBS and transferred into a 1.5 mL tube. The cells were again spun at 1,500 xg for 5 m and the PBS was removed. Each cell pellets was resuspended in 500 μL of lysis buffer (50 mM HEPES pH 7.5, 8 M Urea, 75 mM NaCl) supplemented with EDTA-free cOmplete™ protease inhibitor cocktail (Roche) and PhosSTOP™ phosphatase inhibitor (Roche). The cells were lysed by passing through a 18G needle 10 times, followed by passing through a 22G needle 10 times. The lysates were spun at 13,000 xg for 15 m and frozen in liquid nitrogen for future use.

### Cellular Proteasome Activity Assays

Stable cell lines of HEK293T with dox-inducible expression of PA26 constructs were split into white 96-well plates (Falcon 353296) with 10,000 cells per well in 100 μL CM. Before plating, the cells were washed three times with CM to clear away any residual Trypsin-EDTA (Gibco 25200056) from the passaging process. Each PA26 construct line was plated as three technical triplicates. Cells were treated with 1 μg/mL doxycycline (Sigma-Aldrich D5207) to induce expression of PA26 and incubated at 37°C for 48 h. 100 μL of reagent from either the Chymotrypsin-like, Trypsin-like, or Caspase-like kits of Cell-Based Proteasome-Glo Assays (Promega) were added to each well. Luminescence readings were immediately taken on a Spectramax M5 microplate reader (Molecular Devices; SoftMax Pro 6.5.1) and the data was plotted and statistically analyzed using GraphPad Prism 10.6.1. Luminescence was normalized to DMSO treated cells.

Lysates used for activity assays were prepared as described above. Total protein concentration of lysates was estimated using BCA. Technical triplicate wells were set up on a 384-well black plate (Griener Bio-One, 781209) for each sample containing 50 μg of total protein per well. Fluorogenic activity substrates of either suc-LLVY-amc (AnaSpec, AS-63892), boc-LRR-amc (AdipoGen, AG-CP3-0014), or FAM-LFP (5-FAM-AKVYPYPMEK(QXL520)-NH_2_; AnaSpec) were added to all wells and immediately read using a Spectramax M5 microplate reader (Molecular Devices; SoftMax Pro 6.5.1). Activity was quantified as RFU/min for all samples. Data was analyzed and plotted using GraphPad Prism 10.6.1.

### Quantitative tandem mass tag (TMT) proteomic analysis

Whole proteome analysis was conducted as previously described^48^. Cell pellets were resuspended in 8 M urea lysis buffer (8 M urea, 75 mM NaCl and 50 mM HEPES, pH 7.5) supplemented with EDTA-free protease and phosphatase inhibitors (Roche). Lysates were then clarified by centrifugation (15,700 x g, 15 min at 4 °C), and protein concentrations were determined using a bicinchoninic acid (BCA) assay (Thermo Fisher Scientific). For each sample, 100 µg of total protein was reduced with 5 mM tris(2-carboxyethyl)phosphine (TCEP) for 30 minutes, alkylated with 14 mM iodoacetamide for 30 minutes in the dark, and the reaction was quenched with 10 mM dithiothreitol (DTT) for 15 minutes, all at room temperature (RT). Next, proteins were precipitated using the methanol-chloroform method, and the resulting protein pellets were resuspended in 200 mM EPPS (pH 8.5). Samples were then digested overnight at 37°C with a mixture of Trypsin/LysC (Thermo Fisher Scientific) at a 1:100 (protease:protein) ratio using an orbital shaker set to 1,500 rpm. After digestion, samples were clarified by centrifugation at 15,700 x g for 10 min, and peptide concentration was measured using the Pierce Quantitative Peptide Assay (Thermo Fisher Scientific). Next, tandem mass tagging (TMT) labeling was performed using the TMTpro-16plex reagents (Thermo Fisher Scientific, Lot. No.: YJ372119). Briefly, 25 μg of peptides was adjusted to a concentration of 1 μg/μL with 200 mM EPPS (pH 8.5), followed by the addition of acetonitrile (ACN) to a final concentration of 30%. Next, each sample was labeled with 50 μg of the corresponding TMT reagent and incubated for 1 hr at RT. Labeling reactions were quenched with 0.3% hydroxylamine (Sigma-Aldrich) for 15 minutes at RT. To assess digestion and labeling efficiency and determine the appropriate mixing ratios for equal peptide loading, a ratio check was performed. For this, 1 µL of each sample was mixed and acidified with 5% formic acid (FA) to bring the final concentration of ACN to 3%. The samples were then desalted using C18 Stop and Go Extraction Tips (STAGE-Tip)^49^, dried using a SpeedVac concentrator, and subsequently analyzed on an Orbitrap Exploris 480 mass spectrometer equipped with a FAIMSpro module (Thermo Fisher Scientific). Based on these analyses, mixing volumes were adjusted, samples pooled into a final solution containing 3% ACN and 3% FA, desalted using 50 mg tC18 SepPak cartridges (Waters), and dried in a SpeedVac. Peptides were then resuspended in 10 mM ammonium bicarbonate (NH_4_HCO_3_) and fractionated by basic pH reversed-phase high-performance liquid chromatography (BPRP-HPLC) using an Agilent Zorbax 300 Extended-C18 column (3.5 μm, 4.6 × 250 mm). Mobile phases consisted of Buffer A (5% ACN, 10 mM NH₄HCO₃, pH 8) and Buffer B (90% ACN, 10 mM NH₄HCO₃, pH 8). Fractions were first collected into 96-well plates and then combined into 24 fractions, from which twelve nonadjacent fractions were selected, desalted using the STAGE-Tip method and dried down in a SpeedVac. Peptides were finally resuspended in 0.1% FA for mass spectrometry analysis. Additional sample details are provided in Supplementary Table 6.

All samples were analyzed in an Orbitrap Exploris 480 mass spectrometer operating in high-resolution MS2 (hrMS2) mode (Thermo Fisher Scientific), equipped with a FAIMSpro module and coupled to a Vanquish Neo ultra-high-performance liquid chromatography (UHPLC) system (Thermo Fisher Scientific). Peptides were separated on a 75 μm inner diameter microcapillary column packed with 15 cm of C18 resin (1.7 μm, 120 Å, Aurora Elite™ IonOptics) at a flow rate of 250 nL/min. The mobile phases consisted of Buffer A (0.1% FA in water) and Buffer B (0.1% FA in ACN). Detailed parameters, including gradient length, FAIMS compensation voltages (CV), MS1 resolution, scan range, MS1 maximum injection time, automatic gain control (AGC), isolation window, higher-energy collision dissociation (HCD), MS2 resolution, MS2 maximum injection time, and AGC, are provided in Supplementary Table 7.

### Proteomic data processing and analysis

RAW mass spectra data were converted to mzML format using MSConvert (ProteoWizard software, v3)^50^ and then searched using MSFragger (v4.1)^51^ against a human protein database downloaded from UniProt (retrieved 2024-06-11, containing 20,420 reviewed, canonical entries). In addition, five extra sequences were included in the FASTA file to account for the wildtype PA26 and its four mutants (see Supplementary Table 8). Decoys and common contaminants were also added to the final FASTA file. Precursor ion mass tolerance was set at 20 ppm and product ion tolerance at 0.02 Da. TMTpro labels on lysine residues and peptide N-termini (+304.20715 Da) and carbamidomethylation of cysteine residues (+57.021 Da) were set as static modifications, while oxidation of methionine residues (+15.995 Da) was set as a variable modification. The search was restricted to tryptic-generated peptides, allowing up to two missed cleavage sites. MSFragger output files were processed using MSBooster^52^, Percolator^53^, and ProteinProphet^54^. Resulting PSM lists and protein groups were adjusted and filtered to a 1% false discovery rate (FDR) using Philosopher (v5.1.1)^55^. TMT reporter ion intensities were extracted from the corresponding MS2 scans using IonQuant^56^. The PSM table generated by Philosopher was filtered for quantitation quality: minimum peptide probability was set to 0.9, precursor ion purity ≥ 50%, minimum MS1 intensity 0.05%, and summed MS2 intensity ≥ 5%. PSMs mapping common contaminants were excluded. Lastly, PSMs were aggregated to peptides and proteins by summing the corresponding intensities.

All proteomic analyses were conducted in R (v4.4.1) and RStudio (v2024.12.0). Key packages included dplyr (v1.1.4) and tidyr (v1.3.1) for data manipulation, ggplot2 (v3.5.1) for data visualization, and edgeR (v4.2.2) and limma (v3.60.6) for normalization and data modeling, respectively.

Data were subject to a two-step normalization to account for both sample loading and compositional biases. Normalized intensities were log2-transformed before being incorporated into the limma^57^ framework for differential expression analysis. A separate lineal model was fitted for each protein using lmFit, comparisons of interest were incorporated via makeContrasts and contrasts.fit, and subsequently fed to eBayes. Lastly, topTable was used to calculate the moderated t-statistic and associated p-value for each protein, and those proteins with adjusted p-values < 0.05 (Benjamin-Hochberg [BH] correction to control for the FDR) were considered significant.

Gene ontology (GO) enrichment analyses were performed using enrichGO, with the set of all proteins quantified in the specific experiment acting as background, and org.Hs.eg.db as the human GO (biological process, molecular function and cellular compartment) database (2024-01). Terms with adjusted p-values < 0.05 (BH correction) were considered significant.

Heatmap of the CP/RP/adaptor subunits of the proteasome: The heatmap shows the log2-fold change (log2FC) of each PA26 construct versus the parental HEK293T cell lines. The proteins depicted were selected based on their belonging to the ubiquitin-proteasome system (UPS) class ‘Proteasome and associated proteins’, as defined by the human Proteostasis Network (PN) annotation^58–60^, which encompasses the following groups: ‘Activators and inhibitors’, ‘Adaptors’, ‘Assembly chaperones’, ‘Associated DUBs’, ‘Associated E3 ligase’, ‘Associated non-Ub enzymes’, ‘Core particle subunit’, ‘Modifiers’, ‘Regulators’ and ‘Regulatory particle subunit’.

Catalytic peptide plots: RAW mass spectra were searched using the same software and parameters as specified in the Methods. The only difference lay in the FASTA file, where the sequences for the pro-peptides and mature catalytic subunits of the human proteasome (PSMB5, PSMB6 and PSMB7) were added as separate units, along with the PA26 sequences. Data processing was also performed in a similar manner. Normalized raw values were scaled to a 100 (the sum of intensities of a peptide across all samples summed up to 100) for plotting purposes.

## REFERENCES

1. Smith, D.M., Benaroudj, N. & Goldberg, A. Proteasomes and their associated ATPases: a destructive combination. J Struct Biol 156, 72–83 (2006).

2. Fabre, B. et al. Label-free quantitative proteomics reveals the dynamics of proteasome complexes composition and stoichiometry in a wide range of human cell lines. J Proteome Res 13, 3027–37 (2014).

3. Opoku-Nsiah, K.A. & Gestwicki, J.E. Aim for the core: suitability of the ubiquitin-independent 20S proteasome as a drug target in neurodegeneration. Transl Res 198, 48–57 (2018).

4. Förster, A., Whitby, F.G. & Hill, C.P. The pore of activated 20S proteasomes has an ordered 7-fold symmetric conformation EMBO J. 22, 4356–4364 (2003).

5. Förster, A., Masters, E.I., Whitby, F.G., Robinson, H. & Hill, C.P. The 1.9 A structure of a proteasome-11S activator complex and implications for proteasome-PAN/PA700 interactions. Mol Cell 18, 589–99 (2005).

6. Smith, D.M. et al. Docking of the proteasomal ATPases’ carboxyl termini in the 20S proteasome’s alpha ring opens the gate for substrate entry. Mol Cell 27, 731–44 (2007).

7. Sadre-Bazzaz, K., Whitby, F.G., Robinson, H., Formosa, T. & Hill, C.P. Structure of a Blm10 complex reveals common mechanisms for proteasome binding and gate opening. Mol Cell 37, 728–35 (2010).

8. Rabl, J. et al. Mechanism of gate opening in the 20S proteasome by the proteasomal ATPases. Mol Cell 30, 360–8 (2008).

9. Buel, G.R., Lu, X. & Walters, K.J. Exploiting the Proteasome for Disease Treatment. in Protein Homeostasis in Drug Discovery 135–177 (2022).

10. Cekala, K., Trepczyk, K., Witkowska, J., Jankowska, E. & Wieczerzak, E. Rpt5-Derived Analogs Stimulate Human Proteasome Activity in Cells and Degrade Proteins Forming Toxic Aggregates in Age-Related Diseases. Int J Mol Sci 25(2024).

11. Jones, C.L., Njomen, E., Sjogren, B., Dexheimer, T.S. & Tepe, J.J. Small Molecule Enhancement of 20S Proteasome Activity Targets Intrinsically Disordered Proteins. ACS Chem Biol 12, 2240–2247 (2017).

12. Myers, N. et al. The Disordered Landscape of the 20S Proteasome Substrates Reveals Tight Association with Phase Separated Granules. Proteomics 18, e1800076 (2018).

13. Pepelnjak, M. et al. Systematic identification of 20S proteasome substrates. Mol Syst Biol 20, 403–427 (2024).

14. Anderson, R.T., Bradley, T.A. & Smith, D.M. Hyperactivation of the proteasome in Caenorhabditis elegans protects against proteotoxic stress and extends lifespan. J Biol Chem 298, 102415 (2022).

15. Gillette, T.G., Kumar, B., Thompson, D., Slaughter, C.A. & DeMartino, G.N. Differential roles of the COOH termini of AAA subunits of PA700 (19 S regulator) in asymmetric assembly and activation of the 26 S proteasome. J Biol Chem 283, 31813–22 (2008).

16. Chen, S. et al. Structural basis for dynamic regulation of the human 26S proteasome. Proc Natl Acad Sci U S A 113, 12991–12996 (2016).

17. Huang, X., Luan, B., Wu, J. & Shi, Y. An atomic structure of the human 26S proteasome. Nat Struct Mol Biol 23, 778–85 (2016).

18. Smith, D.M. et al. ATP binding to PAN or the 26S ATPases causes association with the 20S proteasome, gate opening, and translocation of unfolded proteins. Mol Cell 20, 687–98 (2005).

19. Toste Rego, A. & da Fonseca, P.C.A. Characterization of Fully Recombinant Human 20S and 20S-PA200 Proteasome Complexes. Mol Cell 76, 138–147 e5 (2019).

20. Stadtmueller, B.M. et al. Structural models for interactions between the 20S proteasome and its PAN/19S activators. J Biol Chem 285, 13–7 (2010).

21. Chuah, J.J.Y., Rexroad, M.S. & Smith, D.M. High resolution structures define divergent and convergent mechanisms of archaeal proteasome activation. Commun Biol 6, 733 (2023).

22. Kandolf, S. et al. Cryo-EM structure of the plant 26S proteasome. Plant Commun 3, 100310 (2022).

23. Dong, Y. et al. Cryo-EM structures and dynamics of substrate-engaged human 26S proteasome. Nature 565, 49–55 (2019).

24. Kumar, B., Kim, Y.C. & DeMartino, G.N. The C terminus of Rpt3, an ATPase subunit of PA700 (19 S) regulatory complex, is essential for 26 S proteasome assembly but not for activation. J Biol Chem 285, 39523–35 (2010).

25. Opoku-Nsiah, K.A. et al. The YPhi motif defines the structure-activity relationships of human 20S proteasome activators. Nat Commun 13, 1226 (2022).

26. Yao, Y. et al. Structural and functional characterizations of the proteasome-activating protein PA26 from Trypanosoma brucei. J Biol Chem 274, 33921–30 (1999).

27. Holehouse, A.S., Ahad, J., Das, R.K. & Pappu, R.V. CIDER: Classification of Intrinsically Disordered Ensemble Regions. Biophysical Journal 108(2015).

28. Kowarz, E., Loscher, D. & Marschalek, R. Optimized Sleeping Beauty transposons rapidly generate stable transgenic cell lines. Biotechnol J 10, 647–53 (2015).

29. Yazgili, A.S. et al. In-gel proteasome assay to determine the activity, amount, and composition of proteasome complexes from mammalian cells or tissues. STAR Protoc 2, 100526 (2021).

30. Ali, A. et al. Differential regulation of the REGgamma-proteasome pathway by p53/TGF-beta signalling and mutant p53 in cancer cells. Nat Commun 4, 2667 (2013).

31. Li, J. & Rechsteiner, M. Molecular dissection of the 11S REG (PA28) proteasome activators. Biochimie 83, 373–383 (2001).

32. Li, J., Powell, S.R. & Wang, X. Enhancement of proteasome function by PA28α overexpression protects against oxidative stress. FASEB J 25, 883–93 (2011).

33. van der Lee, R. et al. Intrinsically disordered segments affect protein half-life in the cell and during evolution. Cell Rep 8, 1832–1844 (2014).

34. Levine, Z.A., Larini, L., LaPointe, N.E., Feinstein, S.C. & Shea, J.E. Regulation and aggregation of intrinsically disordered peptides. Proc Natl Acad Sci U S A 112, 2758–63 (2015).

35. Liu, Z. et al. An overview of PROTACs: a promising drug discovery paradigm. Mol Biomed 3, 46 (2022).

36. Bekes, M., Langley, D.R. & Crews, C.M. PROTAC targeted protein degraders: the past is prologue. Nat Rev Drug Discov 21, 181–200 (2022).

37. Coleman, R.A. & Trader, D.J. Development and Application of a Sensitive Peptide Reporter to Discover 20S Proteasome Stimulators. ACS Comb Sci 20, 269–276 (2018).

38. Perez-Riverol, Y. et al. The PRIDE database at 20 years: 2025 update. Nucleic Acids Res 53, D543–D553 (2025).

39. Strom, A.R. et al. HP1alpha is a chromatin crosslinker that controls nuclear and mitotic chromosome mechanics. Elife 10(2021).

40. Wang, F. et al. General and robust covalently linked graphene oxide affinity grids for high-resolution cryo-EM. Proc Natl Acad Sci U S A 117, 24269–24273 (2020).

41. Zheng, S.Q. et al. MotionCor2: anisotropic correction of beam-induced motion for improved cryo-electron microscopy. Nat Methods 14, 331–332 (2017).

42. de la Rosa-Trevin, J.M. et al. Scipion: A software framework toward integration, reproducibility and validation in 3D electron microscopy. J Struct Biol 195, 93–9 (2016).

43. Punjani, A., Rubinstein, J.L., Fleet, D.J. & Brubaker, M.A. cryoSPARC: algorithms for rapid unsupervised cryo-EM structure determination. Nat Methods 14, 290–296 (2017).

44. Kimanius, D., Forsberg, B.O., Scheres, S.H. & Lindahl, E. Accelerated cryo-EM structure determination with parallelisation using GPUs in RELION-2. Elife 5(2016).

45. Emsley, P., Lohkamp, B., Scott, W.G. & Cowtan, K. Features and development of Coot. Acta Crystallogr D Biol Crystallogr 66, 486–501 (2010).

46. Croll, T.I. ISOLDE: a physically realistic environment for model building into low-resolution electron-density maps. Acta Crystallogr D Struct Biol 74, 519–530 (2018).

47. Adams, P.D. et al. PHENIX: a comprehensive Python-based system for macromolecular structure solution. Acta Crystallogr D Biol Crystallogr 66, 213–21 (2010).

48. Luo, Q. et al. Targetable leukaemia dependency on noncanonical PI3Kgamma signalling. Nature 630, 198–205 (2024).

49. Rappsilber, J., Ishihama, Y. & Mann, M. Stop and Go Extraction Tips for Matrix-Assisted Laser Desorption/Ionization, Nanoelectrospray, and LC/MS Sample Pretreatment in Proteomics. Anal. Chem. 75, 663–670 (2003).

50. Kessner, D., Chambers, M., Burke, R., Agus, D. & Mallick, P. ProteoWizard: open source software for rapid proteomics tools development. Bioinformatics 24, 2534–6 (2008).

51. Kong, A.T., Leprevost, F.V., Avtonomov, D.M., Mellacheruvu, D. & Nesvizhskii, A.I. MSFragger: ultrafast and comprehensive peptide identification in mass spectrometry-based proteomics. Nat Methods 14, 513–520 (2017).

52. Yang, K.L. et al. MSBooster: improving peptide identification rates using deep learning-based features. Nat Commun 14, 4539 (2023).

53. The, M., MacCoss, M.J., Noble, W.S. & Kall, L. Fast and Accurate Protein False Discovery Rates on Large-Scale Proteomics Data Sets with Percolator 3.0. J Am Soc Mass Spectrom 27, 1719–1727 (2016).

54. Nesvizhskii, A.I., Keller, A., Kolker, E. & Aebersold, R. A Statistical Model for Identifying Proteins by Tandem Mass Spectrometry. Anal. Chem. 75, 4646–4658 (2003).

55. da Viega Leprevost, F., et al. Philosopher: a versatile toolkit for shotgun proteomics data analysis. Nat Methods 17, 869–870 (2020).

56. Yu, F., Haynes, S.E. & Nesvizhskii, A.I. IonQuant Enables Accurate and Sensitive Label-Free Quantification With FDR-Controlled Match-Between-Runs. Mol Cell Proteomics 20, 100077 (2021).

57. Ritchie, M.E. et al. limma powers differential expression analyses for RNA-sequencing and microarray studies. Nucleic Acids Res 43, e47 (2015).

58. Elsasser, S., et al. A Comprehensive Enumeration of the Human Proteostasis Network. 1. Components of Translation, Protein Folding, and Organelle-Specific Systems. bioRxiv (2022).

59. Elsasser, S., et al. A Comprehensive Enumeration of the Human Proteostasis Network. 2. Components of the Autophagy-Lysosome Pathway. bioRxiv (2023).

60. Elsasser, S., et al. Survey of the human proteostasis network: the ubiquitin-proteasome system. bioRxiv (2026).

