## Supplementary Information for "Resolving the Activation Mechanism of the Human 20S Proteasome"

Table of Contents


| | Page |
|---|---|
| **Supplementary Data** | |
| Supplementary Data 1. TMT-MS Protein Quantifications | 2 |
| Supplementary Data 2. GO Annotation Analysis of TMT-MS Protein Hits | 2 |
| **Supplementary Tables** | |
| Supplementary Table 1. Cryo-EM data collection, refinement, and validation statistics. | 3 |
| Supplementary Table 2. Primers Used for Creating PA26 P5 Constructs by Site Directed Mutagenesis. | 5 |
| Supplementary Table 3. Primers Used for Cloning PA26 Constructs into pSBtet-RP Vector. | 5 |
| Supplementary Table 4. Peptides Used in Biochemical Assays | 6 |
| Supplementary Table 5. Antibodies Used in Western Blots | 6 |
| Supplementary Table 6. Sample Details for Quantitative TMT Proteomic Analysis | 7 |
| Supplementary Table 7. Mass Spectrometry Settings | 7 |
| Supplementary Table 8. Additional Sequences Added to FASTA Input File for Quantitative TMT Proteomic Analysis | 8 |
| **Supplemental Videos** | |
| Supplementary Video 1. H20S Proteasome Gate Changes Across PA26-bound Structures | 8 |
| **Supplemental Figures** | |
| Supplementary Figure 1. Proteasome Activation Relies on Sequence Conserved Leucine. | 9 |
| Supplementary Figure 2. The P5 Position Plays an Essential Role in PA26-Mediated Proteasome Activation. | 11 |
| Supplementary Figure 3. Cryo-EM Data Processing and Validation for h20S•$PA26_{P5F}$. | 13 |
| Supplementary Figure 4. Cryo-EM Data Processing and Validation for h20S•$PA26_{P5W}$. | 14 |

Supplementary Figure 5. Cryo-EM Data Processing and Validation for h20S•PA26$_{P5Y}$. 16
Supplementary Figure 6. Cryo-EM Data Processing and Validation for h20S•PA26$_{P5V}$. 17
Supplementary Figure 7. Cryo-EM Data Processing and Validation for h20S•PA26$_{FFT}$. 18
Supplementary Figure 8. P5 Mutations Attenuate 20S Gate Opening. 19
Supplementary Figure 9. P5 Leucine Opens the Gates Through Precise Steric Displacement. 20
Supplementary Figure 10. The P5 Position Dictates Relative Proteasome Activity in Cells. 21

**Supplementary Data**

**Supplementary Data 1. TMT-MS Protein Quantifications**

**Supplementary Data 2. GO Annotation Analysis of TMT-MS Protein Hits**

#### Supplementary Tables

#### Supplementary Table 1. Cryo-EM data collection, refinement, and validation statistics.

| | h20S•PA26$_{P5F}$ | h20S•PA26$_{P5W}$ | h20S•(PA26$_{P5W}$)$_2$ | h20S•PA26$_{P5Y}$ | h20S•PA26$_{P5V}$ |
|---|---|---|---|---|---|
| **PDB accession code** | **12CN** | **12CO** | **12CP** | **12CQ** | **12CR** |
| **EMDB accession code** | **76309** | **76310** | **76311** | **76312** | **76313** |
| **Data collection and processing** | | | | | |
| Microscope and camera | Titan Krios, K3 | Titan Krios, K3 | Titan Krios, K3 | Titan Krios, K3 | Titan Krios, K3 |
| Magnification | 105,000 | 105,000 | 105,000 | 105,000 | 105,000 |
| Voltage (kV) | 300 | 300 | 300 | 300 | 300 |
| Data acquisition software | SerialEM | SerialEM | SerialEM | SerialEM | SerialEM |
| Exposure navigation | Image shift | Image shift | Image shift | Image shift | Image shift |
| Electron exposure ($e^-$/Å$^2$) | 46 | 46 | 46 | 46 | 46 |
| Defocus range (μm) | -0.8 to -1.8 | -0.8 to -1.8 | -0.8 to -1.8 | -0.8 to -1.8 | -0.8 to -1.8 |
| Pixel size (Å) | 0.834 | 0.834 | 0.834 | 0.834 | 0.834 |
| Symmetry imposed | C1 | C1 | C1 | C1 | C1 |
| Initial particle images (no.) | 300,401 | 336,680 | 336,680 | 186,686 | 100,667 |
| Final particle images (no.) | 96,347 | 83,165 | 81,078 | 21,162 | 23,781 |
| Map resolution (Å) | 2.6 | 2.5 | 2.6 | 3.0 | 2.8 |
| FSC threshold | 0.143 | 0.143 | 0.143 | 0.143 | 0.143 |
| Map resolution range (Å) | 2.3-15 | 2.3-15 | 2.3-14 | 2.6-43 | 2.3-46 |
| **Refinement** | | | | | |
| Model resolution (Å) | 2.9 | 2.8 | 3.2 | 3.6 | 3.4 |
| FSC threshold | 0.5 | 0.5 | 0.5 | 0.5 | 0.5 |
| Map sharpening B factor (Å$^2$) | -58.5 | -62.4 | -62.0 | -53.1 | -51.5 |
| **Model composition** | | | | | |
| Non-hydrogen atoms | 58,148 | 57,503 | 68,775 | 57,523 | 57,452 |
| Protein residues | 7,677 | 7,668 | 9,101 | 7,672 | 7,665 |
| Ligands | 0 | 0 | 0 | 0 | 0 |
| **B factors (Å$^2$)** | | | | | |
| Protein | 55.21 | 25.32 | 41.30 | 69.65 | 56.16 |
| Ligands | N/A | N/A | N/A | N/A | N/A |
| **R.M.S. deviations** | | | | | |
| Bond lengths (Å) | 0.004 | 0.004 | 0.005 | 0.004 | 0.004 |
| Bond angles (°) | 1.024 | 1.025 | 1.048 | 1.022 | 1.031 |
| **Validation** | | | | | |
| MolProbity score | 0.99 | 0.83 | 0.97 | 0.98 | 0.85 |
| Clash score | 1.48 | 1.17 | 1.52 | 1.21 | 1.19 |
| Rotamer outliers (%) | 0.12 | 0.07 | 0.09 | 0.05 | 0.04 |
| **Ramachandran plot** | | | | | |
| Favored (%) | 97.51 | 98.07 | 97.67 | 97.23 | 97.93 |
| Allowed (%) | 2.49 | 1.93 | 2.33 | 2.77 | 2.04 |
| Outliers (%) | 0.00 | 0.00 | 0.00 | 0.00 | 0.04 |

#### Supplementary Table 1. (continued)

| | h20S•PA26$_{FFT}$ |
|---|---|
| **PDB accession code** | **12CS** |
| **EMDB accession code** | **76314** |
| **Data collection and processing** | |
| Microscope and camera | Glacios, K2 |
| Magnification | 53,937 |
| Voltage (kV) | 200 |
| Data acquisition software | SerialEM |
| Exposure navigation | Image shift |
| Electron exposure ($e^-$/Å$^2$) | 66 |
| Defocus range (μm) | -0.5 to -2.5 |
| Pixel size (Å) | 0.927 |
| Symmetry imposed | C1 |
| Initial particle images (no.) | 248,905 |
| Final particle images (no.) | 46,362 |
| Map resolution (Å) | 3.1 |
| FSC threshold | 0.143 |
| Map resolution range (Å) | 2.5-10 |
| **Refinement** | |
| Model resolution (Å) | 3.3 |
| FSC threshold | 0.5 |
| Map sharpening B factor (Å$^2$) | -79.8 |
| **Model composition** | |
| Non-hydrogen atoms | 59,467 |
| Protein residues | 7,639 |
| Ligands | 0 |
| **B factors (Å$^2$)** | |
| Protein | 63.27 |
| Ligands | N/A |
| **R.M.S. deviations** | |
| Bond lengths (Å) | 0.004 |
| Bond angles (°) | 1.061 |
| **Validation** | |
| MolProbity score | 0.93 |
| Clash score | 1.75 |
| Rotamer outliers (%) | 0.00 |
| **Ramachandran plot** | |
| Favored (%) | 98.29 |
| Allowed (%) | 1.71 |
| Outliers (%) | 0.00 |

**Supplementary Table 2. Primers Used for Creating PA26 P5 Constructs by Site Directed Mutagenesis.**

| Primer Name | Sequence |
|---|---|
| PA26P5F_for | 5'-GGGAAACTTCTCGTATTACACATAACATTGGAAGTGG-3' |
| PA26P5F_rev | 5'-AATACGAGAAGTTTCCCGTACGAGGCTGAATGAGC-3' |
| PA26P5W_for | 5'-GGGAAACTGGTCGTATTACACATAACATTGGAAGTGG-3' |
| PA26P5W_rev | 5'-AATACGACCAGTTTCCCGTACGAGGCTGAATGAGC-3' |
| PA26P5Y_for | 5'-GGGAAACTACTCGTATTACACATAACATTGGAAGTGG-3' |
| PA26P5Y_rev | 5'-AATACGAGTAGTTTCCCGTACGAGGCTGAATGAGC-3' |
| PA26P5V_for | 5'-GGGAAACGTGTCGTATTACACATAACATTGGAAGTGG-3' |
| PA26P5V_rev | 5'-AATACGACACGTTTCCCGTACGAGGCTGAATGAGC-3' |
| PA26P5I_for | 5'-GGGAAACATCTCGTATTACACATAACATTGGAAGTGG-3' |
| PA26P5I_rev | 5'-AATACGAGATGTTTCCCGTACGAGGCTGAATGAGC-3' |

**Supplementary Table 3. Primers Used for Cloning PA26 Constructs into pSBtet-RP Vector.**

| Primer Name | Sequence |
|---|---|
| PA26_SBfor | 5'-GAAAGGCCTCTGAGGCCACCATGCCACCGAAACGCGCC-3' |
| PA26YYT_SBrev | 3'-GTCCAAACTCATCAATGTATCTTATCATGTCTATCGTTATGTGTAATACGACAGGTTTCCCGTACG-5' |
| PA26YAT_SBrev | 3'-GTCCAAACTCATCAATGTATCTTATCATGTCTATCGTTATGTGGCATACGACAGGTTTCCCGTACG-5' |
| PA26P5F_SBrev | 3'-GTCCAAACTCATCAATGTATCTTATCATGTCTATCGTTATGTGTAATACGAGAAGTTTCCCGTACG-5' |
| PA26P5W_SBrev | 3'-GTCCAAACTCATCAATGTATCTTATCATGTCTATCGTTATGTGTAATACGACCAGTTTCCCGTACG-5' |
| PA26P5Y_SBrev | 3'-GTCCAAACTCATCAATGTATCTTATCATGTCTATCGTTATGTGTAATACGAGTAGTTTCCCGTACG-5' |
| PA26P5V_SBrev | 3'-GTCCAAACTCATCAATGTATCTTATCATGTCTATCGTTATGTGTAATACGACACGTTTCCCGTACG-5' |

**Supplementary Table 4. Peptides Used in Biochemical Assays**

| Peptide Sequence | N-Terminal Modification | C-Terminal Modification |
|---|---|---|
| NLSYYT | Acetylation | None |
| NLSYAT | Acetylation | None |
| NFSYYT | Acetylation | None |
| NWSYYT | Acetylation | None |
| NYSYYT | Acetylation | None |
| NVSYYT | Acetylation | None |
| NISYYT | Acetylation | None |
| N{Nle}SYYT | Acetylation | None |
| N{Nva}SYYT | Acetylation | None |
| NASYYT | Acetylation | None |
| NKSYYT | Acetylation | None |
| NRSYYT | Acetylation | None |
| NHSYYT | Acetylation | None |
| NDSYYT | Acetylation | None |
| NESYYT | Acetylation | None |
| NNSYYT | Acetylation | None |
| NQSYYT | Acetylation | None |
| NSSYYT | Acetylation | None |
| NTSYYT | Acetylation | None |
| NPSYYT | Acetylation | None |
| NMSYYT | Acetylation | None |
| NGSYYT | Acetylation | None |
| NCSYYT | Acetylation | None |
| KLSYYT | Acetylation | None |
| SLSYYT | Acetylation | None |
| TLSYYT | Acetylation | None |
| RLSYYT | Acetylation | None |
| QLSYYT | Acetylation | None |
| VLSYYT | Acetylation | None |
| LLSYYT | Acetylation | None |
| ILSYYT | Acetylation | None |

**Supplementary Table 5. Antibodies Used in Western Blots**

| Name | Catalog # | Manufacturer | Lot/Batch# |
|---|---|---|---|
| anti β-actin polyclonal mouse | A2228 | Sigma | 0000097749 |
| anti PSMA5 polyclonal rabbit | ab11437 | abcam | 1013902-4 |

**Supplementary Table 6. Sample Details for Quantitative TMT Proteomic Analysis**

| TMT kit | TMT-18plex | | |
|---|---|---|---|
| **Channel** | **Sample Information** | **Mutant** | **Replicate** |
| 126 | HEK293T | WT | 1 |
| 127N | HEK293T_PA26_P5V | P5V | 2 |
| 127C | HEK293T | WT | 3 |
| 128N | HEK293T_PA26_YYT | YYT | 1 |
| 128C | HEK293T_PA26_YYT | YYT | 2 |
| 129N | HEK293T_PA26_YYT | YYT | 3 |
| 129C | HEK293T_PA26_YAT | YAT | 1 |
| 130N | HEK293T_PA26_YAT | YAT | 2 |
| 130C | HEK293T_PA26_YAT | YAT | 3 |
| 131N | HEK293T_PA26_P5W | P5W | 1 |
| 131C | HEK293T_PA26_P5W | P5W | 2 |
| 132N | HEK293T_PA26_P5W | P5W | 3 |
| 132C | HEK293T_PA26_P5V | P5V | 1 |
| 133N | HEK293T | WT | 2 |
| 133C | HEK293T_PA26_P5V | P5V | 3 |

**Supplementary Table 7. Mass Spectrometry Settings**

| | |
|---|---|
| Instrument | Exploris 480 |
| MS runs | 12 |
| Method | FAIMS-HR-MS2 |
| Method duration (min) | 140 |
| FAIMS CV | -40/-60/-75 |
| MS1 detector | Orbitrap |
| MS1 Resolution | 120k |
| MS1 Scan Range (m/z) | 400-1600 |
| MS1 Max Inj Time (ms) | Auto |
| MS1 AGC Target | Standard |
| MS2 Iso Window (m/z) | 0.7 |
| HCD Colision Energy (%) | 38 |
| MS2 detector | Orbitrap |
| MS2 Resolution | 30k |

**Supplementary Table 8. Additional Sequences Added to FASTA Input File for Quantitative TMT Proteomic Analysis**

| Protein Name | FASTA Sequence |
|---|---|
| PA26$_{E102A}$ | MPPKRAALIQNLRDSYTETSSFAVIEEWAAGTLQEIEGIAKAAAEA<br>HGVIRNSTYGRAQAEKSPEQLLGVLQRYQDLCHNVYCQAETIRTV<br>IAIRIPEHKEADNLGVAVQHAVLKIIDELEIKTLGSGEKSGSGGAPTP<br>IGMYALREYLSARSTVEDKLLGSVDAESGKTKGGSQSPSLLLELR<br>QIDADFMLKVELATTHLSTMVRAVINAYLLNWKKLIQPRTGSDHMV<br>S |
| PA26$_{YYT}$ | MPPKRAALIQNLRDSYTETSSFAVIEEWAAGTLQEIEGIAKAAAEA<br>HGVIRNSTYGRAQAEKSPEQLLGVLQRYQDLCHNVYCQAETIRTV<br>IAIRIPEHKEADNLGVAVQHAVLKIIDELEIKTLGSGEKSGSGGAPTP<br>IGMYALREYLSARSTVEDKLLGSVDAESGKTKGGSQSPSLLLELR<br>QIDADFMLKVELATTHLSTMVRAVINAYLLNWKKLIQPRTGNLSYYT |
| PA26$_{P5W}$ | MPPKRAALIQNLRDSYTETSSFAVIEEWAAGTLQEIEGIAKAAAEA<br>HGVIRNSTYGRAQAEKSPEQLLGVLQRYQDLCHNVYCQAETIRTV<br>IAIRIPEHKEADNLGVAVQHAVLKIIDELEIKTLGSGEKSGSGGAPTP<br>IGMYALREYLSARSTVEDKLLGSVDAESGKTKGGSQSPSLLLELR<br>QIDADFMLKVELATTHLSTMVRAVINAYLLNWKKLIQPRTGNWSYY<br>T |
| PA26$_{P5V}$ | MPPKRAALIQNLRDSYTETSSFAVIEEWAAGTLQEIEGIAKAAAEA<br>HGVIRNSTYGRAQAEKSPEQLLGVLQRYQDLCHNVYCQAETIRTV<br>IAIRIPEHKEADNLGVAVQHAVLKIIDELEIKTLGSGEKSGSGGAPTP<br>IGMYALREYLSARSTVEDKLLGSVDAESGKTKGGSQSPSLLLELR<br>QIDADFMLKVELATTHLSTMVRAVINAYLLNWKKLIQPRTGNVSYY<br>T |
| PA26$_{YAT}$ | MPPKRAALIQNLRDSYTETSSFAVIEEWAAGTLQEIEGIAKAAAEA<br>HGVIRNSTYGRAQAEKSPEQLLGVLQRYQDLCHNVYCQAETIRTV<br>IAIRIPEHKEADNLGVAVQHAVLKIIDELEIKTLGSGEKSGSGGAPTP<br>IGMYALREYLSARSTVEDKLLGSVDAESGKTKGGSQSPSLLLELR<br>QIDADFMLKVELATTHLSTMVRAVINAYLLNWKKLIQPRTGNLSYAT |

#### Supplementary Videos

**Supplementary Video 1. h20S Proteasome Gate Changes Across PA26-bound Structures.** Top view of h20S is shown with $\alpha$1 (cyan), $\alpha$2 (green), $\alpha$3 (yellow), $\alpha$4 (orange), $\alpha$5 (red), $\alpha$6 (purple), and $\alpha$7 (blue). The video cycles across gate opening intermediates and states starting with the closed apo-structure (PDB: 6RGQ) and following the order PA26$_{P5V}$ (PDB: 12CR), PA26$_{P5Y}$ (PDB: 12CQ), PA26$_{P5W}$ (PDB: 12CP), PA26$_{P5F}$ (PDB: 12CN), and PA26$_{YYT}$ (PDB: 6XMJ).

#### Supplementary Figures

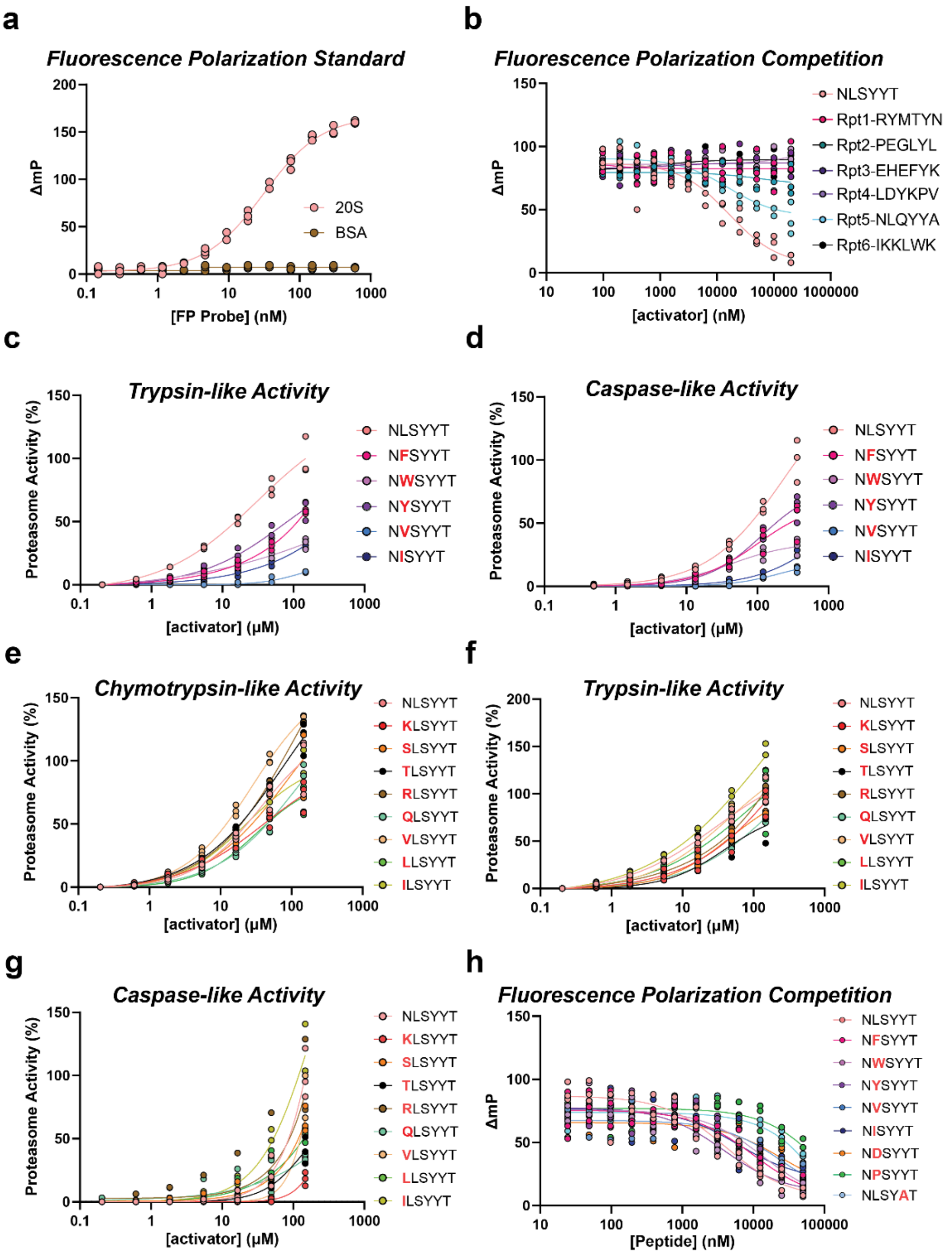


**Supplementary Figure 1. Proteasome Activation Relies on Sequence Conserved Leucine. a)** Fluorescence polarization standard curve of the FP probe binding to h20S proteasome. The FP probe binds to 20S with $EC_{50}$ = 31.15 ± 3.34 nM. Average curve fits are shown with data plotted individually (n=3). **b)** Fluorescence polarization competition assay of peptides against the FP probe used in panel (a). Average curve fits are shown with data plotted individually (n=3). $K_{I,app}$ was calculated from each curve, converted to $K_I$, and reported in Figure 1e. **c)** Proteasome trypsin-like activity assay with peptides derived from NLSYYT with P5 substitutions (red letters). Maximum activity was normalized to NLSYYT. Average curve fits are shown with data plotted individually (n=3). **d)** Proteasome caspase-like activity assay with peptides derived from NLSYYT with P5 substitutions. Maximum activity was normalized to NLSYYT. Average curve fits are shown with data plotted individually (n=3). **e)** Proteasome chymotrypsin-like activity assay with peptides derived from NLSYYT with P6 substitutions. Maximum activity was normalized to NLSYYT. Average curve fits are shown with data plotted individually (n=3). P6 substitutions include KLSYYT (red), SLSYYT (orange), TLSYYT (black), RLSYYT (brown), QLSYYT (aqua), VLSYYT (beige), LLSYT (green), ILSYYT (gold). **f)** Proteasome trypsin-like activity assay with peptides derived from NLSYYT with P6 substitutions. Maximum activity was normalized to NLSYYT. Average curve fits are shown with data plotted individually (n=3) **g)** Proteasome caspase-like activity assay with peptides derived from NLSYYT with P6 substitutions. Maximum activity was normalized to NLSYYT. Average curve fits are shown with data plotted individually (n=3). **h)** Fluorescence polarization competition assay with peptides derived from NLSYYT with P5 substitutions against the FP probe used in panel (a). Average curve fits are shown with data plotted individually (n=3). $K_{I,app}$ was determined from each curve, converted to $K_I$, and reported with error in Figure 1h.

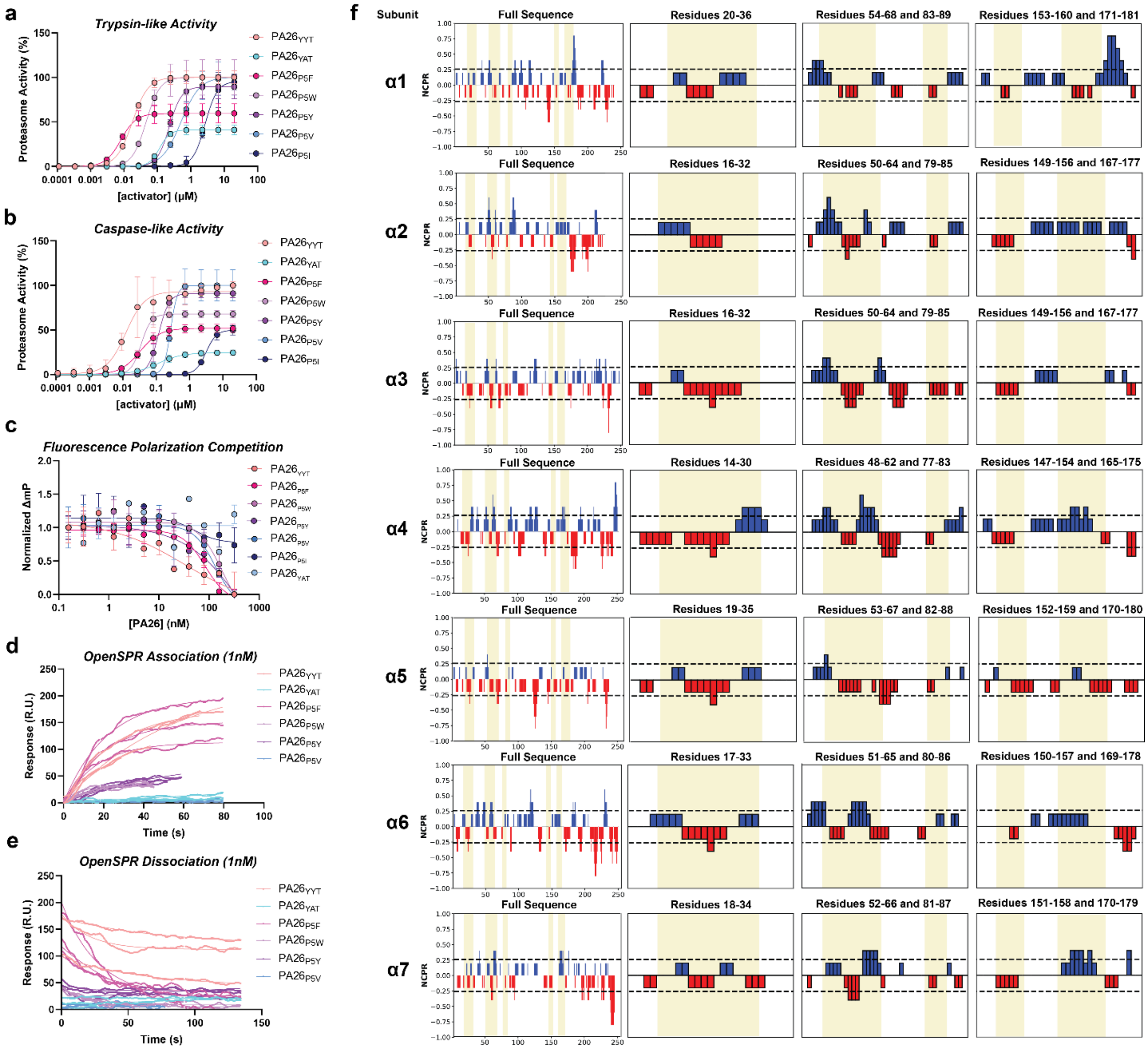


**Supplementary Figure 2. The P5 Position Plays an Essential Role in PA26-Mediated Proteasome Activation. a)** Proteasome trypsin-like activity assay with PA26 constructs: $PA26_{YYT}$ ($EC_{50}$ = 17.91 ± 0.35 nM), $PA26_{P5F}$ ($EC_{50}$ = 8.09 ± 1.30 nM), $PA26_{P5W}$ ($EC_{50}$ = 227.5 ± 18.30 nM), $PA26_{P5Y}$ ($EC_{50}$ = 45.71 ± 8.54 nM), $PA26_{P5V}$ ($EC_{50}$ = 507.6 ± 65.11 nM), and $PA26_{P5I}$ ($EC_{50}$ = 2964 ± 957.3 nM). Maximum activity was normalized to $PA26_{YYT}$. Average curve fits are shown with data plotted individually (n=3). **b)** Proteasome caspase-like activity assay with PA26 constructs: $PA26_{YYT}$ ($EC_{50}$ = 9.60 ± 0.25 nM), $PA26_{P5F}$ ($EC_{50}$ = 33.47 ± 4.21 nM), $PA26_{P5W}$ ($EC_{50}$ = 119.9 ± 17.90 nM), $PA26_{P5Y}$ ($EC_{50}$ = 33.77 ± 5.92 nM), $PA26_{P5V}$ ($EC_{50}$ = 259.2 ± 1.83 nM), and $PA26_{P5I}$ ($EC_{50}$ = 3111 ± 400.1 nM). Maximum activity was normalized to $PA26_{YYT}$. Average curve fits are shown with data plotted individually (n=3). **c)** Fluorescence polarization competition assay comparing proteasome binding of PA26 constructs against the FP probe used in Figure S1a. Average curve fits are shown with data plotted individually (n=3). $K_{I,app}$ was calculated from each curve, converted to $K_I$, and reported in Figure 2e. **d)** OpenSPR association curves for 1 nM of each PA26 construct injected onto immobilized 20S. Individual curve fits are shown with data plotted individually (n=3). $k_{on}$ was calculated for each curve and reported in the data table in Figure 2f. **e)** OpenSPR dissociation curves for 1 nM of each PA26 construct injected onto immobilized 20S. Individual curve fits are shown with data plotted individually (n=3) $k_{off}$ was calculated for each curve and reported in the data table in Figure 2f. **f)** Net charge per residue calculations for each 20S $\alpha$-subunit were calculated using local CIDER[27]. Specific surfaces within the PA binding pockets are highlighted within each subunit and shown as individual plots to highlight energetic differences between pockets.

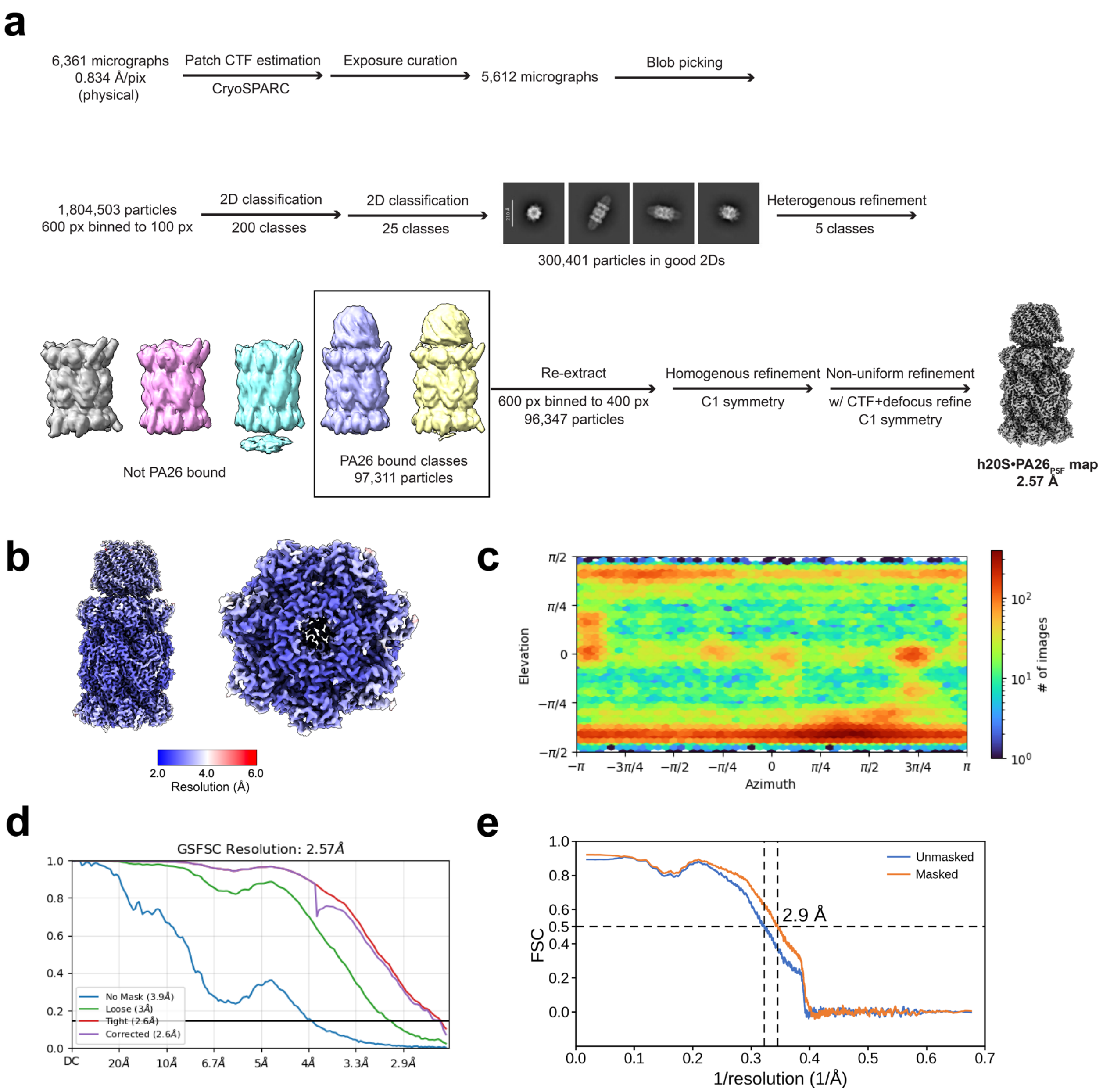


**Supplementary Figure 3. Cryo-EM Data Processing and Validation for h20S•PA26$_{P5F}$. a)** Cryo-EM data processing flowchart for the h20S•PA26$_{P5F}$ dataset. **b)** Local resolution estimation of finalized map. The right panel shows a top view of the 20S core particle at the PA26-bound interface. **c)** Viewing direction distributions for finalized particle stack. **d)** Fourier Shell Correlation (FSC) curves between refined half-maps. **e)** Map-to model FSC curves. Displayed model resolution was determined using the masked map.

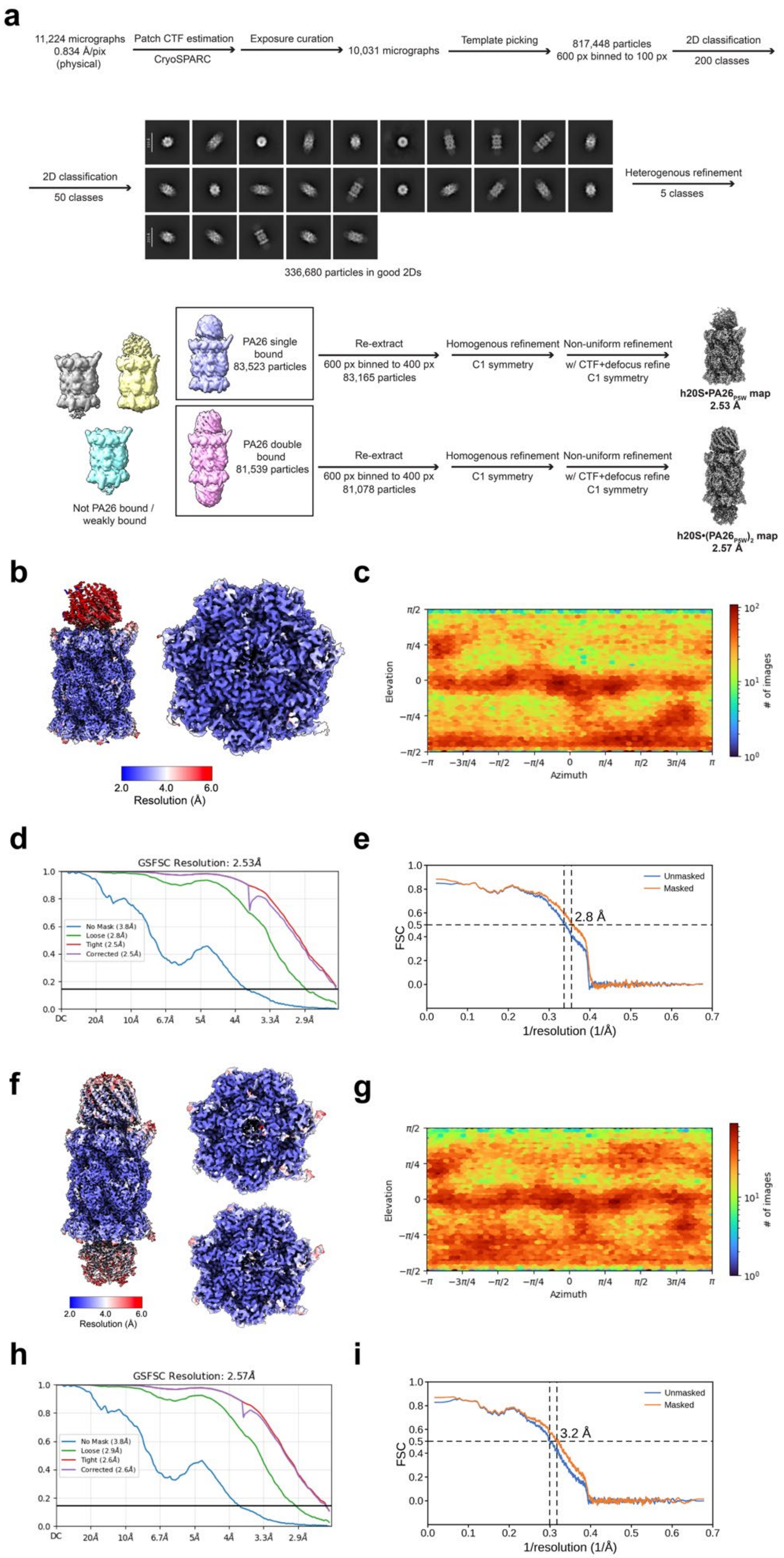
a
11,224 micrographs
0.834 Å/pix
(physical)
Patch CTF estimation
CryoSPARC
Exposure curation
10,031 micrographs
Template picking
817,448 particles
600 px binned to 100 px
2D classification
200 classes
2D classification
50 classes
Heterogenous refinement
5 classes
336,680 particles in good 2Ds
PA26 single
bound
83,523 particles
Re-extract
600 px binned to 400 px
83,165 particles
Homogenous refinement
C1 symmetry
Non-uniform refinement
w/ CTF+defocus refine
C1 symmetry
h20S•PA26PSW map
2.53 Å
PA26 double
bound
81,539 particles
Re-extract
600 px binned to 400 px
81,078 particles
Homogenous refinement
C1 symmetry
Non-uniform refinement
w/ CTF+defocus refine
C1 symmetry
Not PA26 bound /
weakly bound
h20S•(PA26PSW)2 map
2.57 Å
b
2.0
4.0
6.0
Resolution (Å)
c
Elevation
Azimuth
### of images
d
GSFSC Resolution: 2.53Å
No Mask (3.8Å)
Loose (2.8Å)
Tight (2.5Å)
Corrected (2.5Å)
e
Unmasked
Masked
2.8 Å
FSC
1/resolution (1/Å)
f
2.0
4.0
6.0
Resolution (Å)
g
Elevation
Azimuth
### of images
h
GSFSC Resolution: 2.57Å
No Mask (3.8Å)
Loose (2.9Å)
Tight (2.6Å)
Corrected (2.6Å)
i
Unmasked
Masked
3.2 Å
FSC
1/resolution (1/Å)

**Supplementary Figure 4. Cryo-EM Data Processing and Validation for h20S•PA26$_{P5W}$.** **a)** Cryo-EM data processing flowchart for the h20S•(PA26$_{P5W}$)$_2$ and h20S•(PA26$_{P5W}$)$_1$ datasets **b)** Local resolution estimation of finalized map for h20S•(PA26$_{P5W}$)$_1$. The right panel shows a top view of the 20S core particle at the PA26-bound interface. **c)** Viewing direction distributions for finalized particle stack for h20S•(PA26$_{P5W}$)$_1$. **d)** Fourier Shell Correlation (FSC) curves between refined half-maps for h20S•(PA26$_{P5W}$)$_1$. **e)** Map-to model FSC curves for h20S•(PA26$_{P5W}$)$_1$. Displayed model resolution was determined using the masked map. **f)** Local resolution estimation of finalized map for h20S•(PA26$_{P5W}$)$_2$. The right panels show top and bottom views of the 20S core particle at the PA26-bound interfaces. **g)** Viewing direction distributions for finalized particle stack for h20S•(PA26$_{P5W}$)$_2$. **h)** Fourier Shell Correlation (FSC) curves between refined half-maps for h20S•(PA26$_{P5W}$)$_2$. **i)** Map-to model FSC curves for h20S•(PA26$_{P5W}$)$_2$. Displayed model resolution was determined using the masked map.

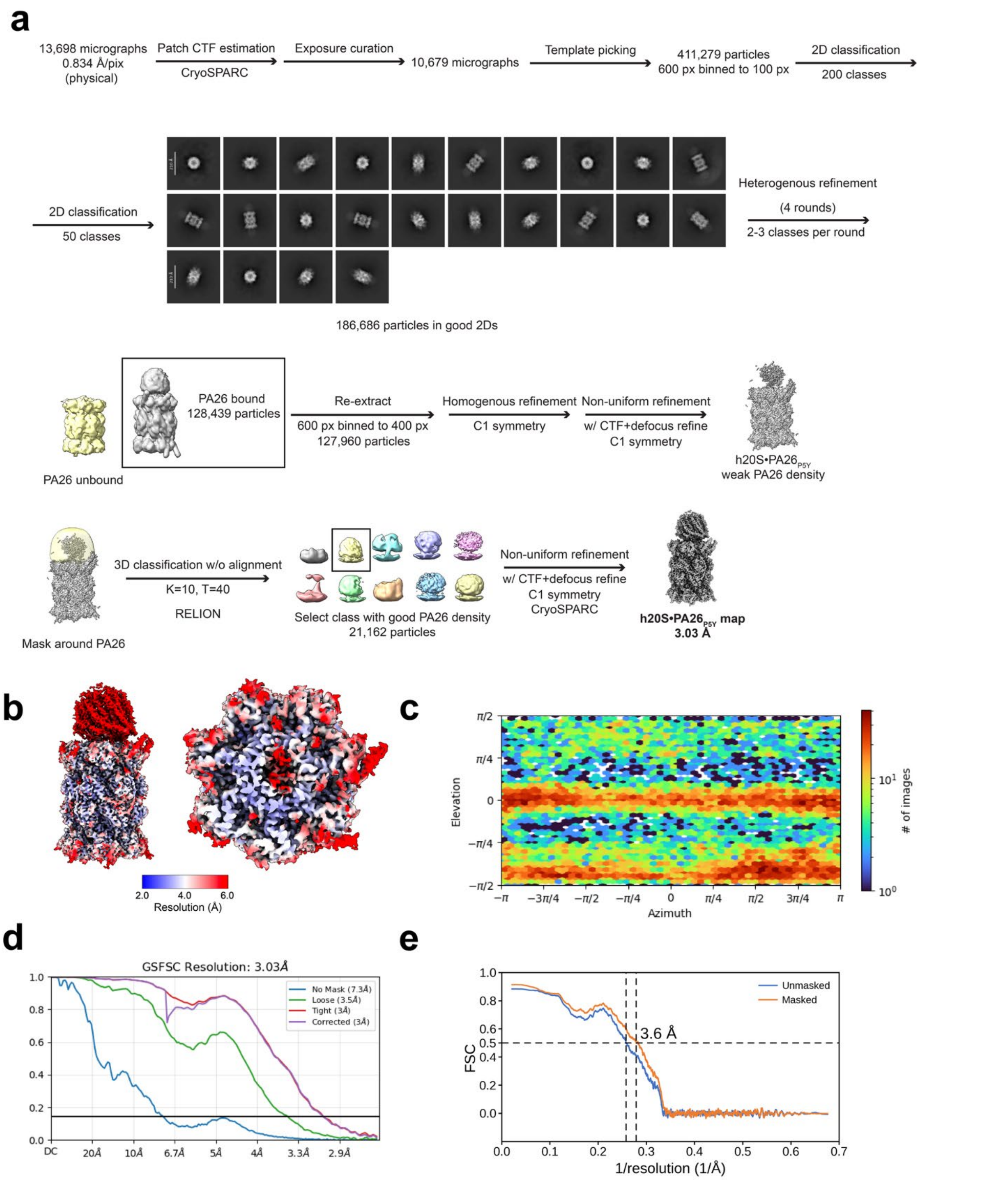


**Supplementary Figure 5. Cryo-EM Data Processing and Validation for h20S•PA26$_{P5Y}$.** **a)** Cryo-EM data processing flowchart for the h20S•PA26$_{P5Y}$ dataset. **b)** Local resolution estimation of finalized map. The right panel shows a top view of the 20S core particle at the PA26-bound interface. **c)** Viewing direction distributions for finalized particle stack. **d)** Fourier Shell Correlation (FSC) curves between refined half-maps. **e)** Map-to model FSC curves. Displayed model resolution was determined using the masked map.

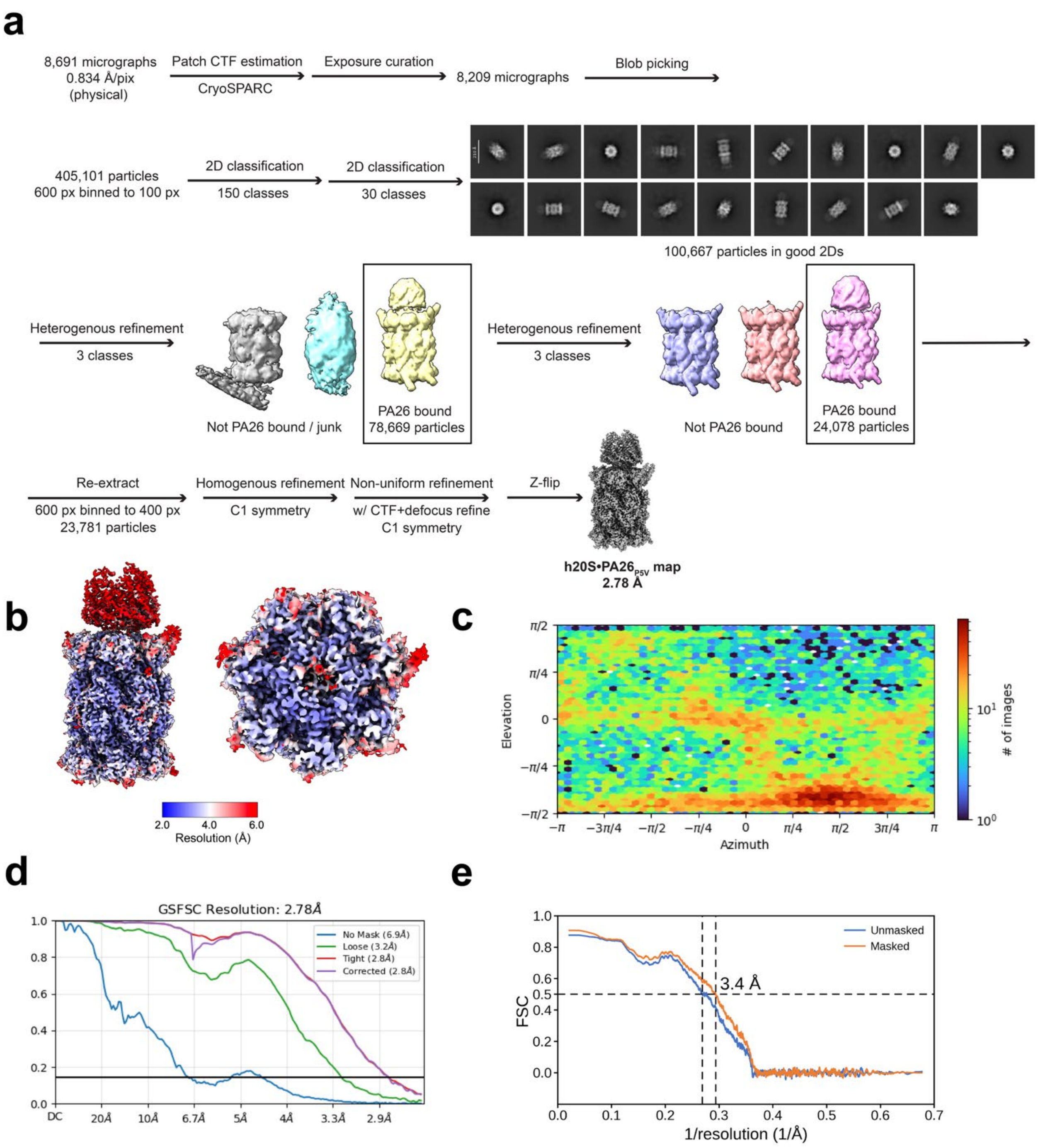


**Supplementary Figure 6. Cryo-EM Data Processing and Validation for h20S•PA26$_{P5V}$.** **a)** Cryo-EM data processing flowchart for the h20S•PA26$_{P5V}$ dataset. **b)** Local resolution estimation of finalized map. The right panel shows a top view of the 20S core particle at the PA26-bound interface. **c)** Viewing direction distributions for finalized particle stack. **d)** Fourier Shell Correlation (FSC) curves between refined half-maps. **e)** Map-to model FSC curves. Displayed model resolution was determined using the masked map.

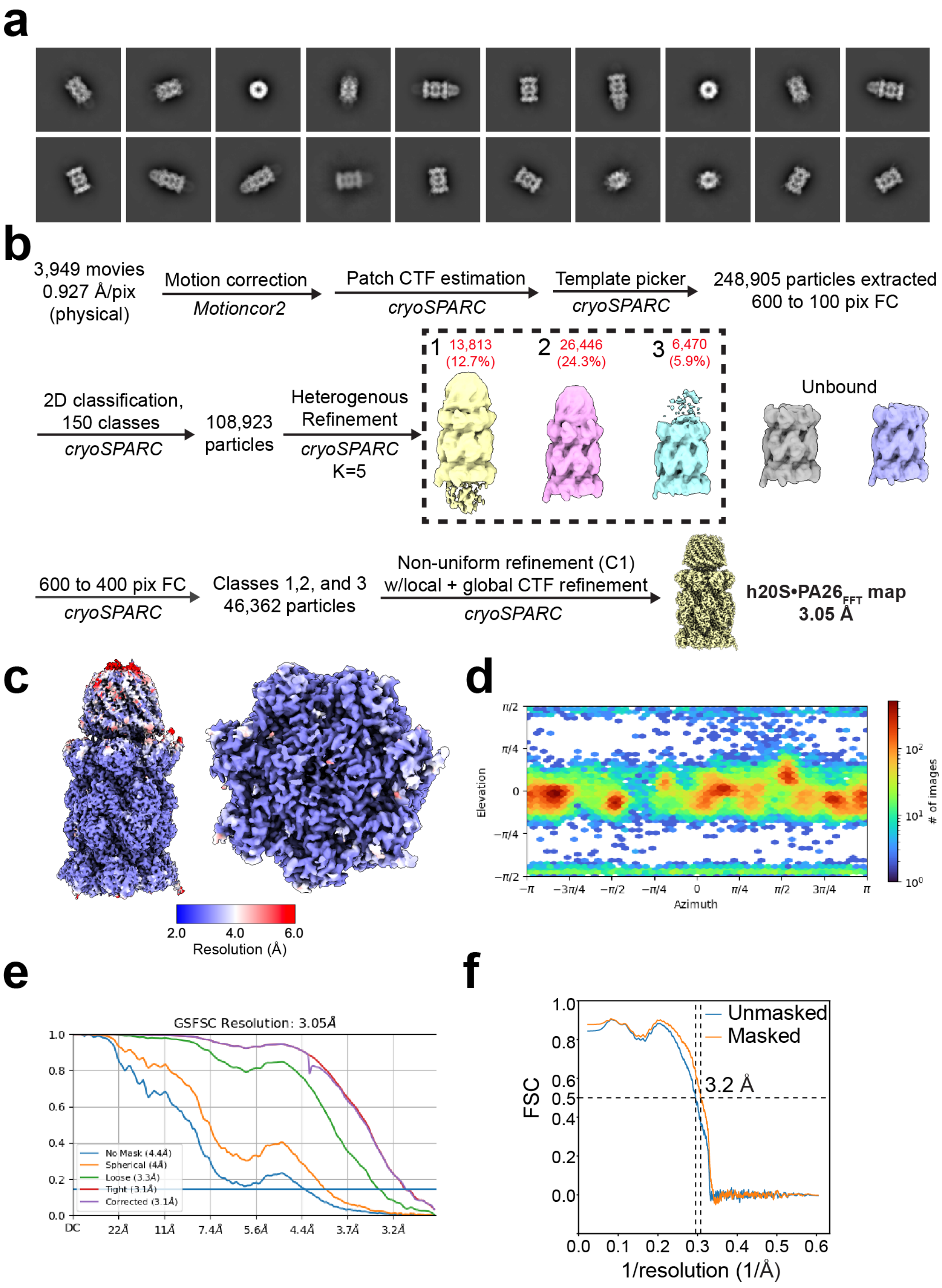


**Supplementary Figure 7. Cryo-EM Data Processing and Validation for h20S•PA26$_{FFT}$. a)** Reference-free 2D class averages of the particles. **b)** Cryo-EM data processing flowchart for the h20S•PA26$_{FFT}$ dataset. **b)** Local resolution estimation of finalized map. The right panel shows a top view of the 20S core particle at the PA26-bound interface. **c)** Viewing direction distributions for finalized particle stack. **d)** Fourier Shell Correlation (FSC) curves between refined half-maps. **e)** Map-to model FSC curves. Displayed model resolution was determined using the masked map.

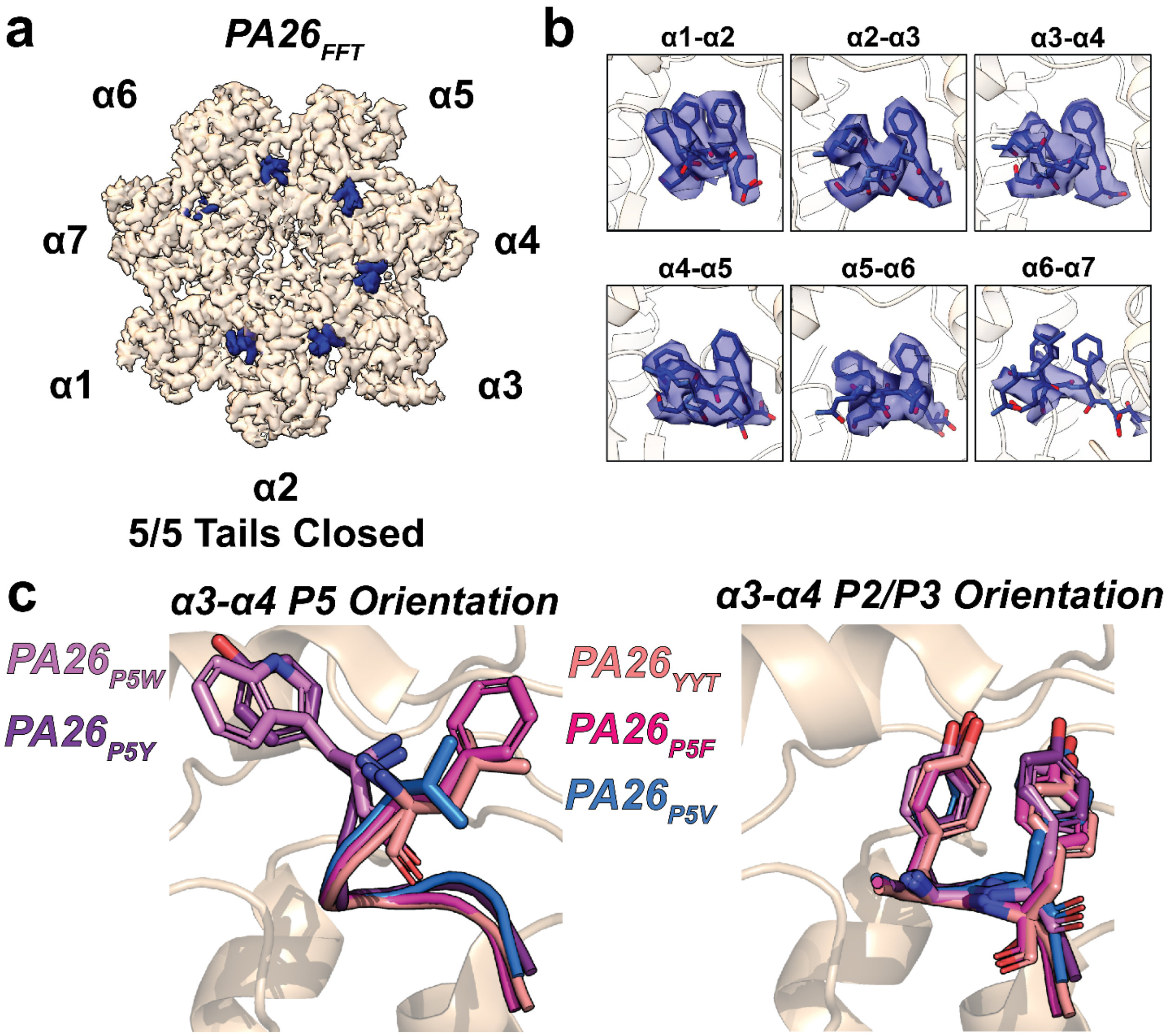


**Supplementary Figure 8. P5 Mutations Attenuate 20S Gate Opening. a)** Top view of cryo-EM density map of h20S bound to $PA26_{FFT}$ (dark blue, PDB: 12CS). PA tails are resolved in $\alpha$-subunit pockets where present. Each $\alpha$-subunit is marked with the yeast homolog nomenclature for simplicity. **b)** Cryo-EM density maps of $PA26_{FFT}$ tails shown in panel (a). Each PA tail density is shown in a grid indicating the structure and corresponding α-subunit pocket. **c)** Cartoon and stick representation of PA26 tails bound to the h20S $\alpha$3-$\alpha$4 pocket with highlighted P5 side chains (left) and P2+P3 side chains (right). While all P2+P3 positions are the same across PA26 tails, the P5 side chain position will vary depending on the identity.

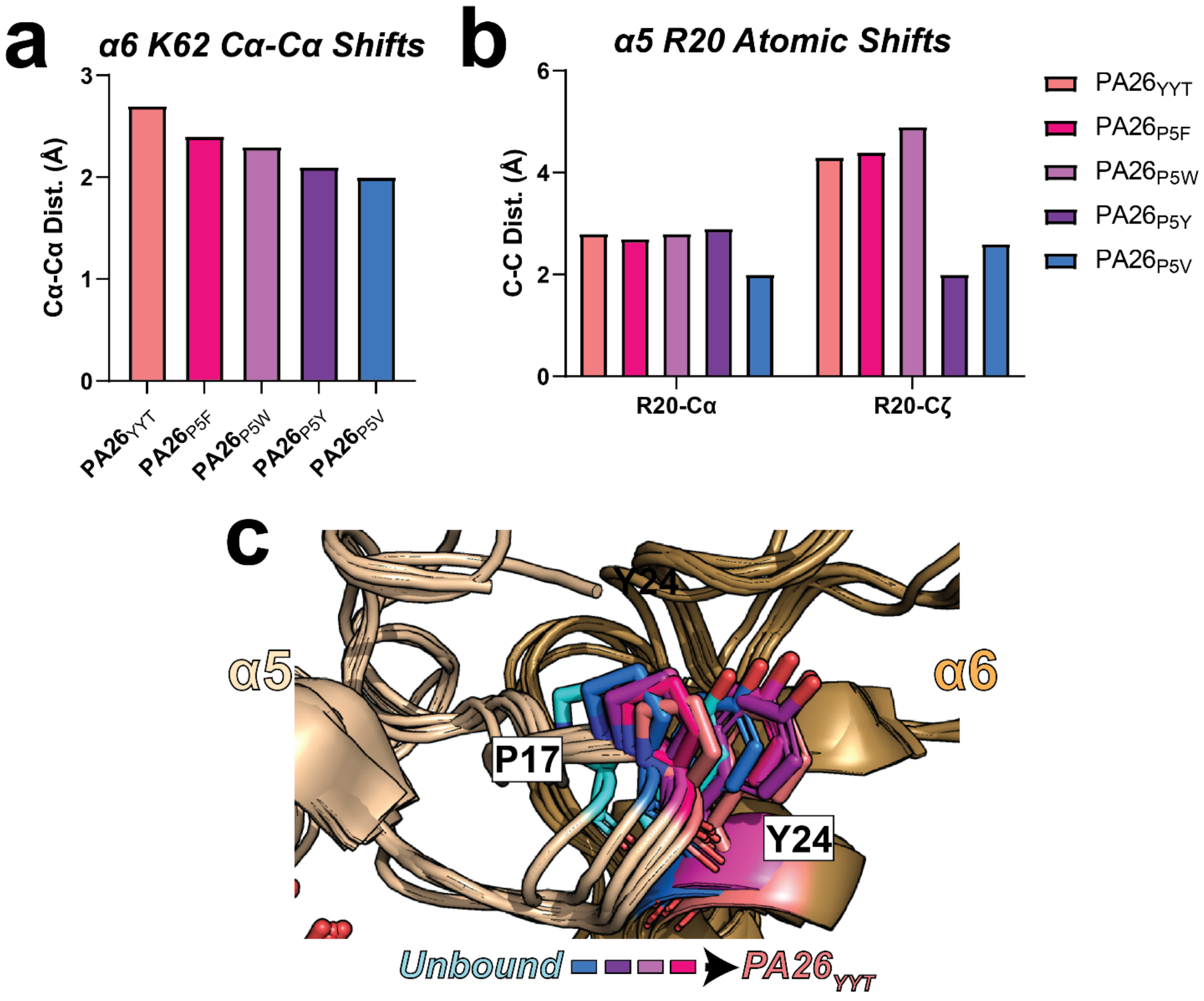


**Supplementary Figure 9. P5 Leucine Opens the Gates Through Precise Steric Displacement. a)** Cα-Cα shifts of K62 in the 20S α6 subunit when bound to $PA26_{YYT}$, $PA26_{P5F}$, $PA26_{P5W}$, $PA26_{P5Y}$, and $PA26_{P5V}$ relative to the unbound 20S structure. **b)** Carbon shifts of backbone Cα and side chain Cζ of R20 in the 20S α5 subunit when bound to $PA26_{YYT}$, $PA26_{P5F}$, $PA26_{P5W}$, $PA26_{P5Y}$, and $PA26_{P5V}$. **c)** Cartoon representation of PA26-bound structures highlighting the positional shift of P17 on the 20S α5 subunit and Y24 on the 20S α6 subunit when bound to $PA26_{YYT}$, $PA26_{P5F}$, $PA26_{P5W}$, $PA26_{P5Y}$, and $PA26_{P5V}$.

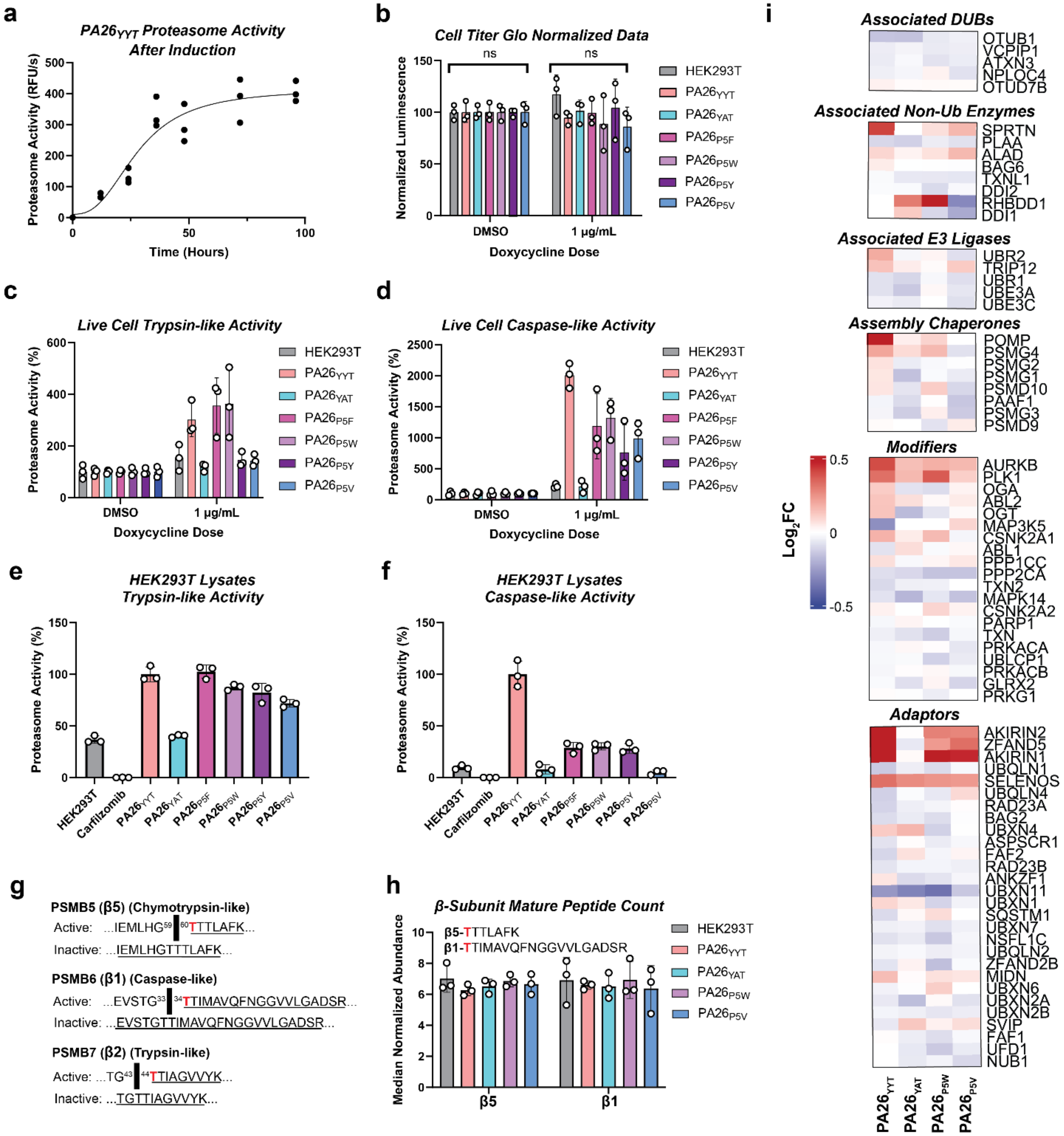


**Supplementary Figure 10. The P5 Position Dictates Relative Proteasome Activity in Cells. a)** Proteasome chymotrypsin-like activity assay where activity was measured at different time points in cells with doxycycline-induced $PA26_{YYT}$ expression. Average curve fit shown with data reported individually as biological replicates (n=3). **b)** Cell Titer Glo fluorescence of different PA26 expressing cell lines treated with either DMSO or 1 µg/mL doxycycline. Data were normalized to DMSO controls. Data reported individually as biological replicates (n=3). **c)** Live cell proteasome trypsin-like activity assay. Activity was normalized to DMSO-treated cells for all cell lines, and activity was measured for cells treated with 1 μg/mL doxycycline. Data reported individually as biological replicates (n=3). Significance values were calculated by two-way ANOVA and compared to $PA26_{YYT}$ where ns = not significant, p<0.1 = *, p<0.01 = **, p<0.001 = ***, and p<0.0001 = ****. **d)** Live cell proteasome caspase-like activity assay. Activity was normalized to DMSO-treated cells for all cell lines, and activity was measured for cells treated with 1 μg/mL doxycycline. Data reported individually as biological replicates (n=3). Significance values were calculated by two-way ANOVA and compared to $PA26_{YYT}$ where ns = not significant, p<0.1 = *, p<0.01 = **, p<0.001 = ***, and p<0.0001 = ****. **e)** Proteasome trypsin-like activity assay on HEK293T lysates. Activity was normalized to $PA26_{YYT}$ for comparison. Data reported individually as biological replicates (n=3). Significance values were calculated by one-way ANOVA and compared to $PA26_{YYT}$ where ns = not significant, p<0.1 = *, p<0.01 = **, p<0.001 = ***, and p<0.0001 = ****. **f)** Proteasome caspase-like activity assay on HEK293T lysates. Activity was normalized to $PA26_{YYT}$ for comparison. Data reported individually as biological replicates (n=3). Significance values were calculated by one-way ANOVA and compared to $PA26_{YYT}$ where ns = not significant, p<0.1 = *, p<0.01 = **, p<0.001 = ***, and p<0.0001 = ****. **g)** Human PSMB5, PSMB6, and PSMB7 peptide sequences as they appear before and after maturation. Yeast homologs are included next to each human subunit name. Black bars show the activation cleavage site, and the underlined sequences are produced upon trypsin digestion of the proteins. The catalytic threonine in each subunit is highlighted in red to show that activity can only occur upon cleavage. **h)** Abundance of peptides that correspond to activated β5 and activated β1 across biological triplicates of each cell line analyzed by TMT as shown in Figure 5e. Data reported individually as biological replicates (n=3). **i)** Heat maps of relative peptide abundance for proteins associated with the UPS or UIPS across the cell lines expressing $PA26_{YYT}$, $PA26_{P5W}$, $PA26_{P5V}$, and $PA26_{YAT}$. These include associated deubiquitinases (DUBs), associated enzymes, associated E3 ligases, proteasome assembly chaperones, proteasome modifying proteins, and proteasome adaptor proteins. Abundance is shown as $Log_2FC$ on a scale of -0.5 (blue) to 0.5 (red). Reported hits are consistent across biological replicates (n=3).
